# Lichen speciation is sparked by a substrate requirement shift and reproduction mode differentiation

**DOI:** 10.1101/2022.04.05.487118

**Authors:** Annina Kantelinen, Christian Printzen, Péter Poczai, Leena Myllys

**Affiliations:** Botany Unit, Finnish Museum of Natural History, P.O. Box 7, FI-00014 University of Helsinki, Finland; Senckenberg Forschungsinstitut und Naturmuseum Frankfurt, Senckenberganlage 25, 60325 Frankfurt am Main, Germany

**Author notes:** Author for correspondence: Annina Kantelinen.

## Abstract

We show that obligate lignicoles in lichenized *Micarea* are predominately asexual whereas most facultative lignicoles reproduce sexually. Our 3 loci phylogenetic analyses (ITS, mtSSU, *Mcm*7) together with ancestral state reconstruction show that the shift in reproduction mode has evolved independently several times within the group and that facultative and obligate lignicoles are sister species. The analyses support the assumption that the ancestor of these species was a facultative lignicole. We hypotezise that a shift in substrate requirement from bark to wood leads to differentiation in reproduction mode and becomes a driver of speciation. This is the first example of lichenized fungi where reproduction mode is linked to substrate requirement. This is also the first example where such a linkage is demonstrated to spark lichen speciation. Our main hypothesis is that obligate species on dead wood need to colonize new suitable substrata relatively fast and asexual reproduction is more effective a strategy for successful colonization. Our main hypothesis for explaining the discovered phenomenon involves the species’ life cycle: species on decaying wood face a significant challenge because the ecological properties of their substratum change relatively fast. When this happens, species need to colonize new suitable substrata. This may set a time limit, where asexual reproduction is a faster and more effective strategy for successful colonization.

## Introduction

Despite increased knowledge on lichen diversity, the factors influencing species richness and speciation are still largely unknown. Existing studies have mostly focused on extrinsic factors and found that diversification events are usually correlated with climatic changes such as climatic cooling events during the Tertiary (Printzen & Lumbsch 2000), aridification during the Oligocene–Miocene transition (Kraichak et al. 2015) and Pleistocene glacial cycles (Leavitt et al. 2013). Only few studies on lichens have considered extrinsic environmental factors and intrinsic lineage-specific traits jointly: Innovations in secondary chemistry (= extracellular products) together with a shift in substrate requirement were found to trigger adaptive radiation in the lichen family Teloschistaceae (Gaya et al. 2015). Increased nitrogen availability after acquisition of cyanobacterial symbionts led to an adaptive radiation in *Placopsis* (L.) Linds (Schneider et al. 2016). Green-algal or cyanobacterial symbiont interactions through time and space may influence diversification in the genus *Sticta* (Schreb.) Ach. (Widhelm et al. 2018). Interplay between intrinsic traits related to reproduction and extrinsic traits related to ecological opportunities are often correlated with shifts in species diversification in other organisms (Vamosi & Vamosi 2011, Wagner et al. 2012, Karunarathne et al. 2018, Nakov et al. 2018) but to our knowledge, this has never been examined in lichenized fungi.

Lichenized fungi have developed diverse reproduction strategies. Many have the ability to reproduce both sexually (ascospores) and asexually (conidia, thallus fragments i.e. soredia, isidia, goniocysts), while some are either sexual or asexual (Tripp 2016, Tripp & Lendemer 2018). Diverse reproduction strategies are at least partly related to lichen symbiosis: asexual reproduction via thallus fragments ensures the continuation of symbiosis (Bowler & Rundell 1975), whereas successful sexual reproduction via ascospores requires that the germinating mycelium makes contact with a compatible free-living photobiont before the lichen thallus can be developed (Buschbom & Mueller 2006). One exception is the asexual propagules produced by the fungal partner, called conidia: they do not contain the photobiont, so they need to find a suitable one from the new substratum they land on. However, both modes of asexual reproduction consume less energy than sexual reproduction and do not rely on the availability of suitable mating partners. Therefore, asexual lichen lineages have been suggested to be more efficient and rapid at colonizing newly exposed substrates (Poelt 1963, Stofer et al. 2006, Ludwig et al. 2017). Recent studies on lichens have shown that asexual lineages are long-lived evolutionarily and can give rise to sexual lineages (i.e. are not evolutionary dead ends), suggesting that asexual lineages may represent a source for evolutionary innovation (see Tripp 2016).

The microlichen genus *Micarea* is an excellent model for studying the effects of reproductive traits and environmental factors on speciation. The genus is widespread worldwide and has lately received much scientific interest, resulting in over 20 new species descriptions (Czarnota 2007; Czarnota & Guzow-Krzemińska 2010; Sérusiaux et al. 2010; van den Boom et al. 2017; Guzow-Krzemińska et al. 2016 & 2019; Kantvilas & Coppins 2019; Launis & Myllys 2019; Launis et al. 2019a, b; van den Boom et al. 2020; Kantelinen et al. 2021a). *Micarea* shows intricate variation in substrate requirements and reproduction modes (Coppins 2009). Certain species are generalists able to grow on various substrata, while some are specialized and living in strict microhabitats (Coppins 1983, Czarnota 2007, Launis & Myllys 2014, Myllys & Launis 2018). A wide range of sexual and asexual propagules is found in *Micarea* (Fig.1), including ascospores, three types of conidia (micro-, meso-, and macroconidia), and thallus fragments called goniocysts that likely act as asexual propagules including both symbiotic partners. Some of the species are predominately sexual, while some frequently lack sexual fruiting structures but bear numerous pycnidia where asexual conidia are produced. The actual roles of the three types of conidia present are not thoroughly known, but Coppins (1983) suggested that mesoconidia act as asexual propagules, based on an observation that several species frequently occur without apothecia but have numerous mesoconidia-producing pycnidia.

**Fig. 1.**
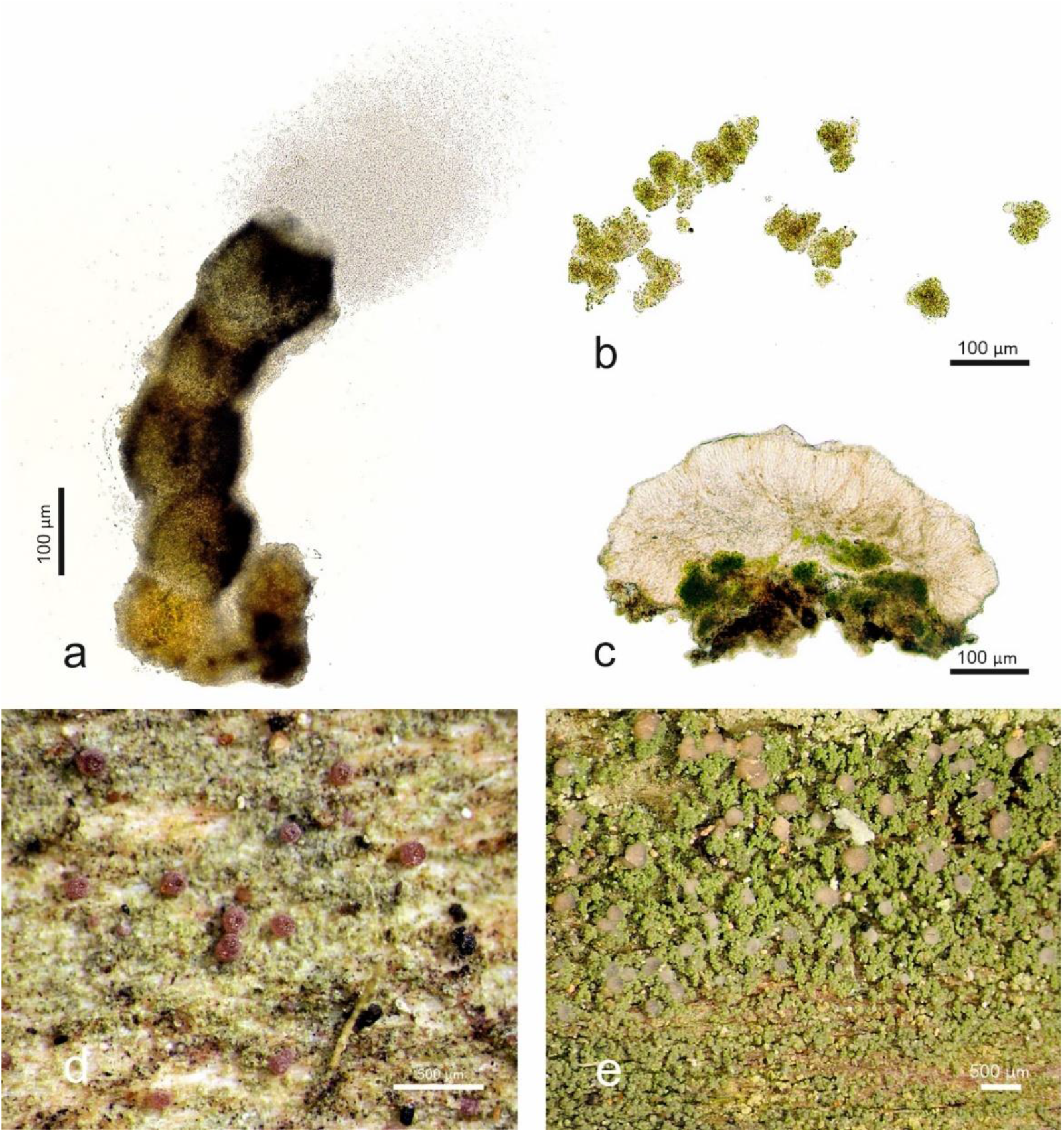
Sexual and asexual structures and reproduction strategies in the *Micarea prasina* group. a.) A pycnidium of *Micarea fennica* extruding asexual mesoconidia (Kantelinen 3220 holotype, H). Mesoconidia are small, likely easily carried by wind and insects and allow long-distance dispersal, b.) Thallus goniocysts of *M. hedlundii* including a mycobiont and a photobiont (Kantelinen 67119, H). Goniocysts are asexual vegetative structures that are relatively big and therefore probably more effective on short distance colonization, c.) Apothecial section of *M. microareolata* (Pykälä 47787, H). Sexual ascospores developed in apothecia are small, likely easily carried by wind and insects, and their development requires a mating partner and more energy than asexual diaspores, d.) Pycnidia and thallus of *M. tomentosa* (Kantelinen 29151, H), e.) Apothecia and thallus of *M. prasina* (Kantelinen 229106, H). Photos A. Kantelinen.

*Micarea* represents one of the most important microlichen genera occurring on dead wood (Spribille et al. 2008). Of the 27 European species in the *M. prasina* group (a monophyletic “core group” including the type species), 17 are facultative lignicoles, five are obligate lignicoles, and four have never been found on dead wood (Czarnota 2007; Spribille et al. 2008; Guzow-Kremínska et al. 2016; Launis et al. 2019a, b; Launis & Myllys 2019; van den Boom et al. 2020). Of the facultative lignicoles, some species are often encountered on dead wood while others rarely occupy the substratum. Furthermore, some of the obligate lignicoles are rare and have very narrow ecological amplitudes, occuring only on wood of specific decay stages (Czarnota 2007; Myllys & Launis 2018; Launis & Myllys 2019).

In this study, the aim is to examine the reproduction modes and evolution of substrate preferences in the *M. prasina* group and how these features affect speciation. The focus is especially on asexual mesopycnidia and presence/absence of sexual apothecia.

## Results

Altogether 516 herbarium specimens were studied. Each specimen was identified to species level, and the reproduction mode and substrate was recorded (Appendix). Our results confirm previous findings that obligate and facultative preference for dead wood are species-specific traits (e.g. Coppins 1983; Czarnota 2007; Launis et al. 2019a&b).

Representative specimens of each taxa were selected for phylogenetic reconstruction (Table 1). The analyses included 3 loci (ITS, *Mcm*7, mtSSU) and consisted of 110 sequences and of 1655 characters. The topologies of the Bayesian and maximum likelihood analyses did not show any strongly supported conflicts, and therefore only the tree obtained from the Bayesian analysis is shown (Fig. 2).

**Table 1.**
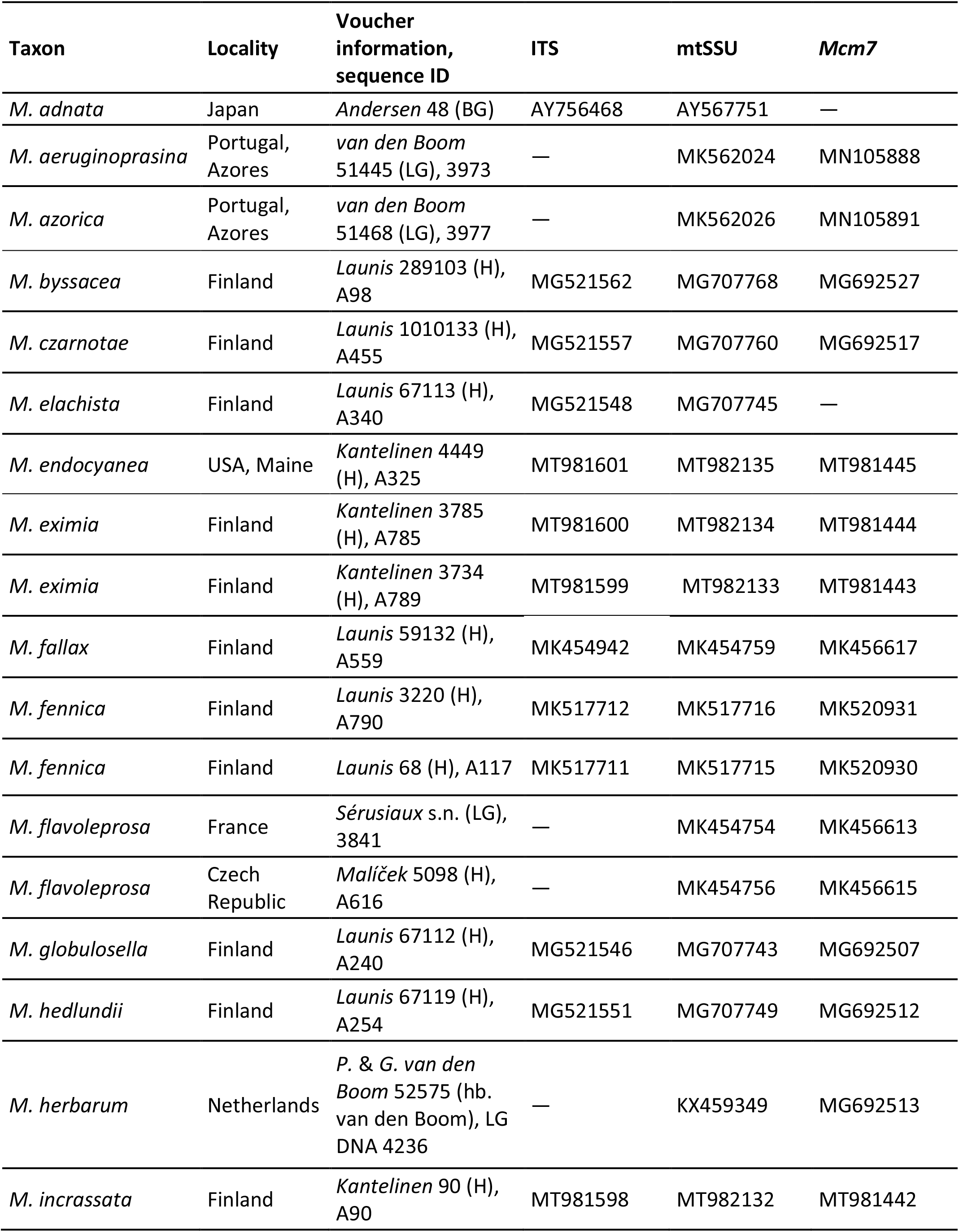
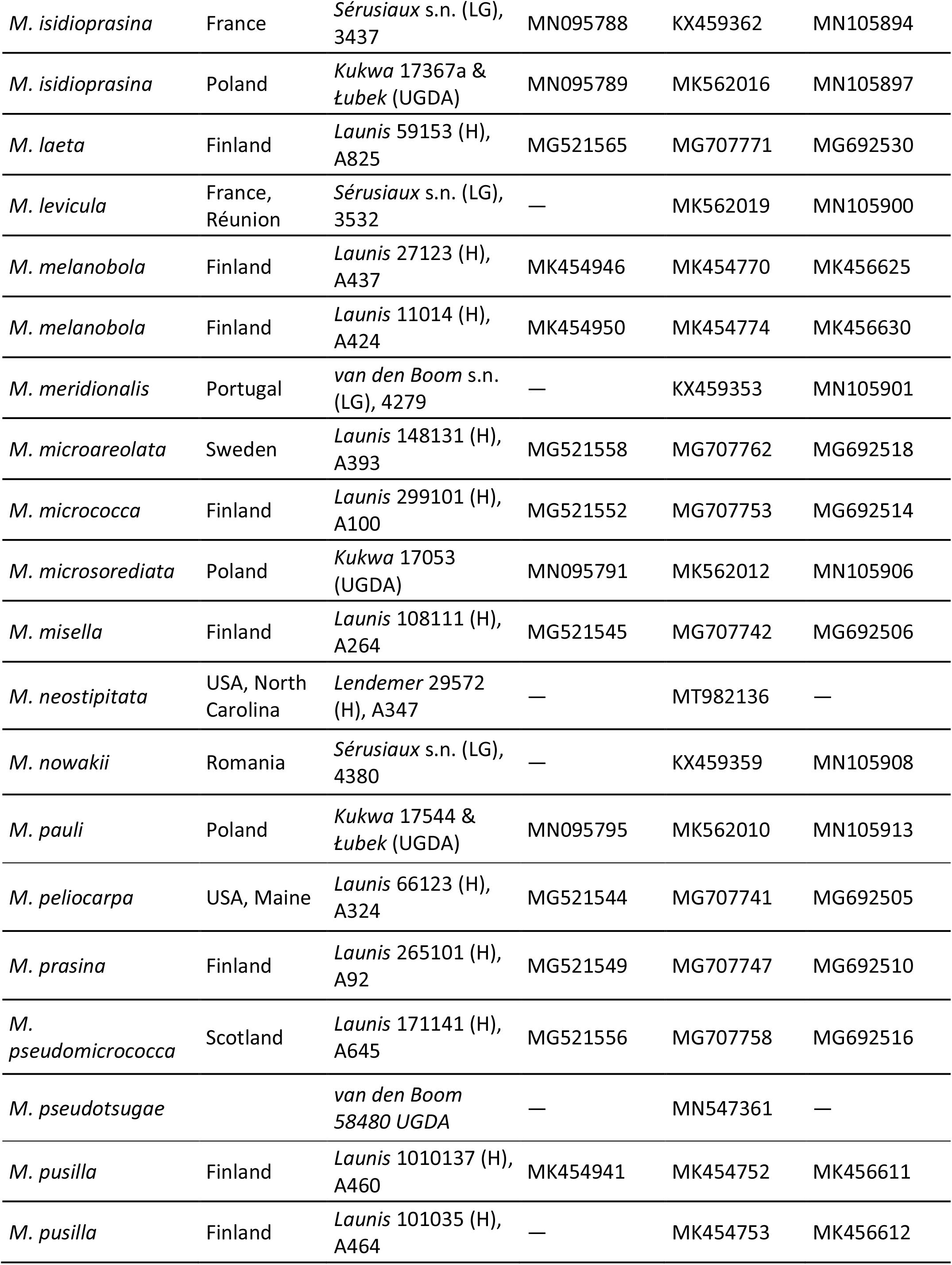
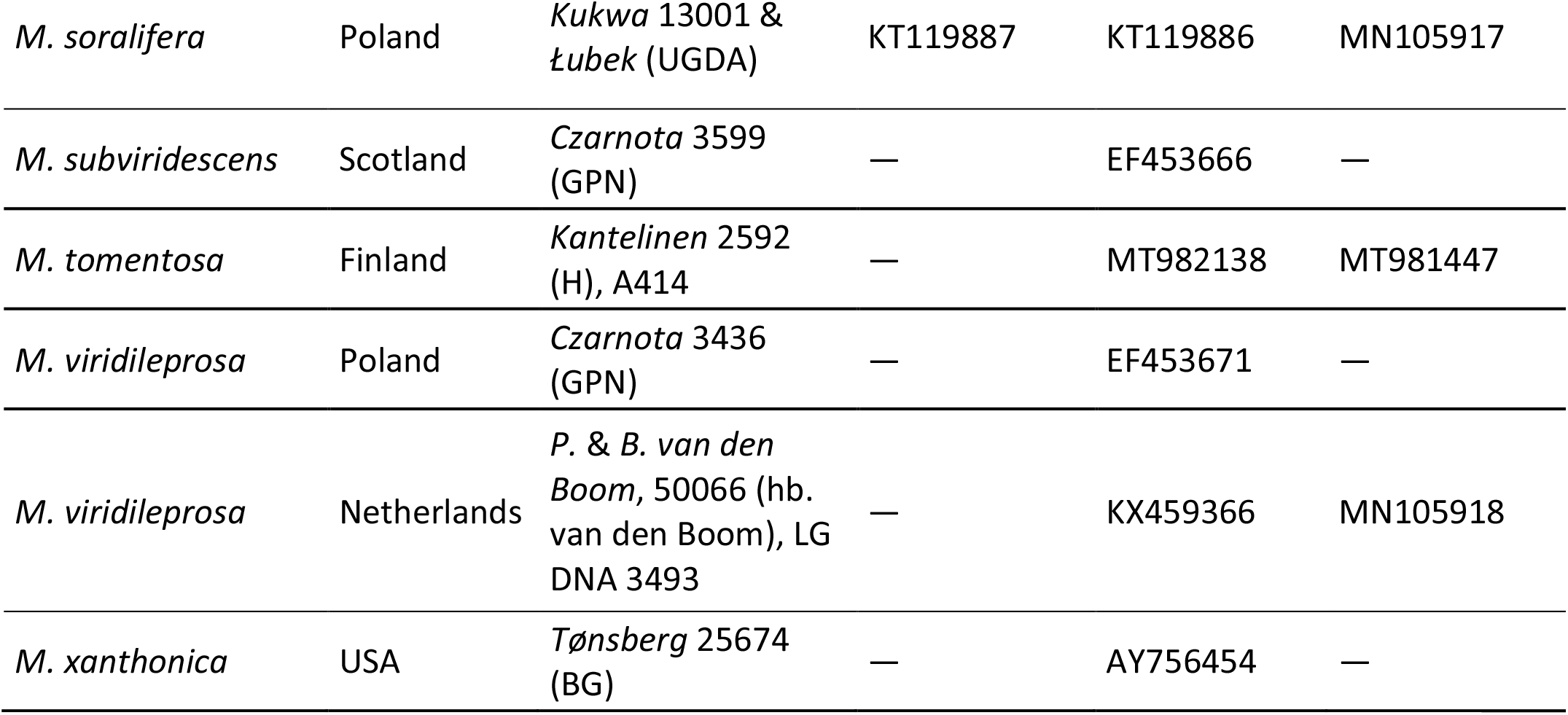
List of *Micarea* specimens used in the phylogenetic analyses with locality, voucher information and GenBank accession numbers.

**Fig. 2.**
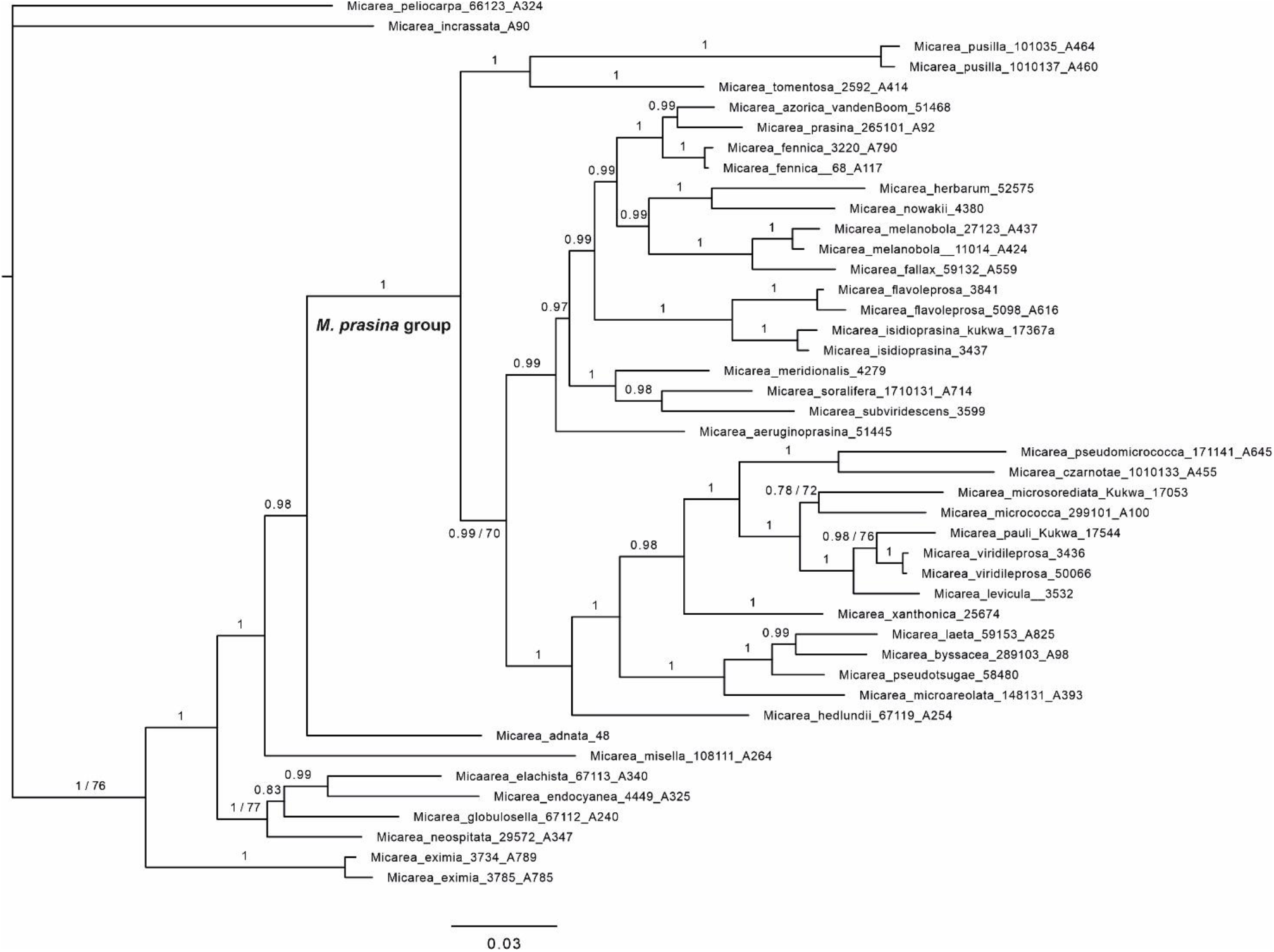
Bayesian tree based on concatenated sequences of ITS, mtSSU and *Mcm7*. Bayesian posterior probabilities are indicated above the nearest branches. Maximum likelihood values are marked if less than 80.

Our results show that obligate lignicoles occupying mid- to late decay stages are predominately asexual while most facultative lignicoles reproduce sexually. Our ancestral state reconstruction shows that the shift in reproduction mode has evolved independently several times within the group and that facultative and obligate lignicoles are sister species (Fig. 3). Furthermore, the ancestral state reconstructions support the ancestor of these species being a facultative lignicole. The correlation between reproductive mode and substrate preference is statistically significant (Table 2).

**Table 2.** Significance of the association between species’ reproduction modes and substrate preferences studied by Fisher Exact Test. The test statistic value is 0.0114, and the result is significant at p < .05.

|  | Sexual | Asexual | Marginal Rows Totals |
| --- | --- | --- | --- |
| Obligate lignicole | 1 | 4 | 5 |
| Facultative lignicole | 14 | 2 | 16 |
| Marginal Column Totals | 15 | 6 | 21 (grand total) |

**Fig. 3.**
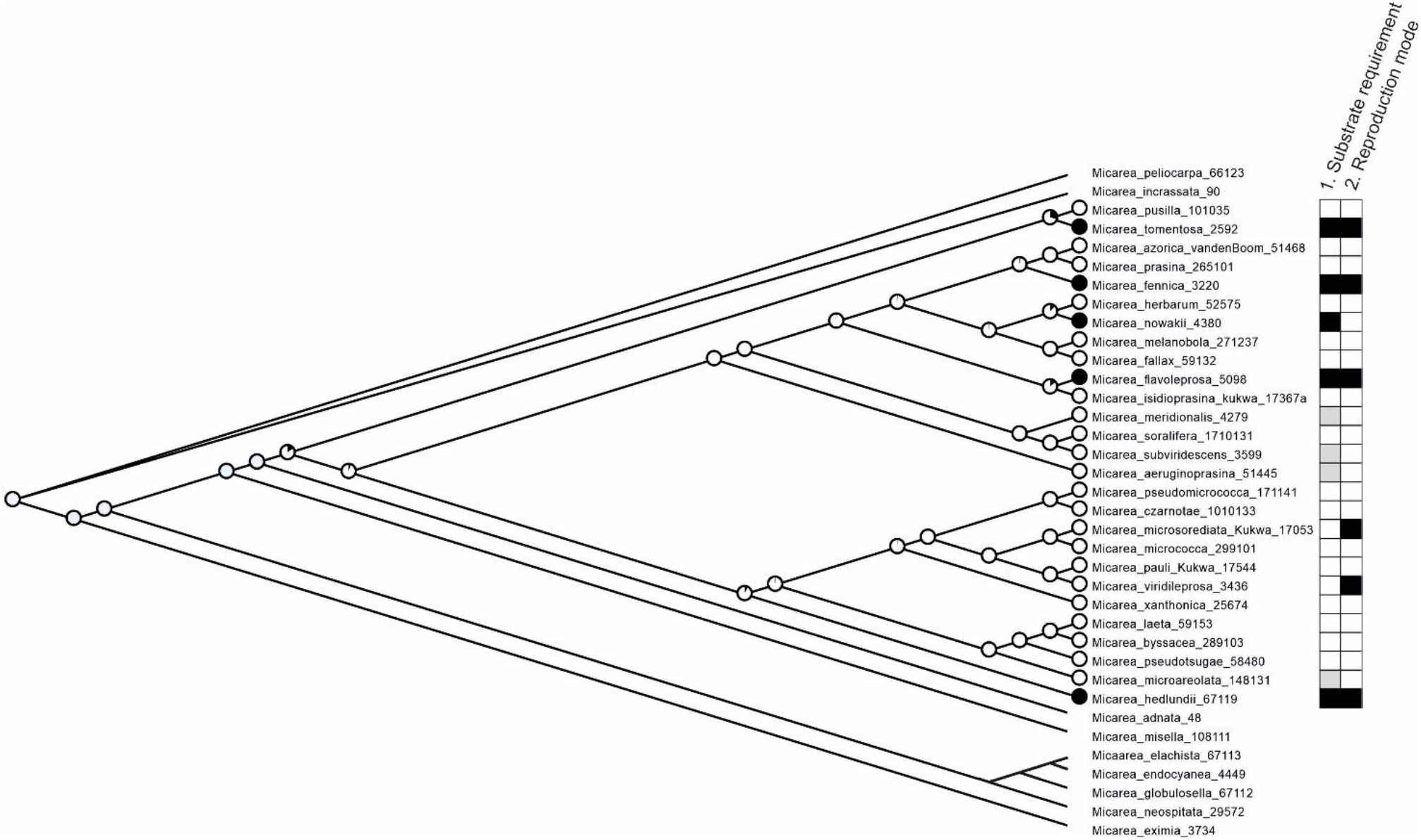
A maximum likelihood phylogram depicting ancestral character state reconstruction of the evolution of obligate lignicoles. Individuals of the same species were pruned and collapsed at the branches of the corresponding nodes. Pies represent probabilities of each ancestor being in two potential states for obligate lignicole (yes = black, no = white). In addition, substratum requirement and reproduction mode are mapped with black, grey and white boxes at the tips of the tree as follows: 1. Substratum requirement: black = obligate lignicole; white = facultative lignicole; grey = never found on dead wood. 2. Reproduction mode: black = predominately asexual; white = predominately sexual.

With strong support values, our phylogenetic reconstruction shows that lignicolous substratum preference has evolved several times independently within the group. Three out of the five obligate lignicoles are nested in the *M. prasina* -complex, i.e. *M. fennica, M. flavoleprosa*, and *M. nowakii. Micarea hedlundii* is resolved to be a separate lineage as a sister of the *M. byssacea* and *M. micrococca* complexes. *Micarea tomentosa* is resolved as a sister of the *M. byssacea, M. micrococca*, and *M. prasina* complexes. The 18 facultative lignicoles in our dataset are found in several lineages within the phylogeny.

## Discussion

Here, we report the *Micarea prasina* group as a first example of lichenized fungi where reproduction mode is linked to substrate requirement (Table 2 and 3). This is also the first example where such a linkage is demonstrated to be a driver of lichen speciation. Our study reveals that the shift in reproduction mode has evolved independently several times within the group and that facultative and obligate lignicoles are sister species (Fig. 3). Before our study, intrinsic and extrinsic factors involved in lichen speciation have rarely been studied, and relationships between lichen substrate requirements and reproduction modes have not been deeply understood (but see Spribille et al. 2008; Resl et al. 2018).

**Table 3.**
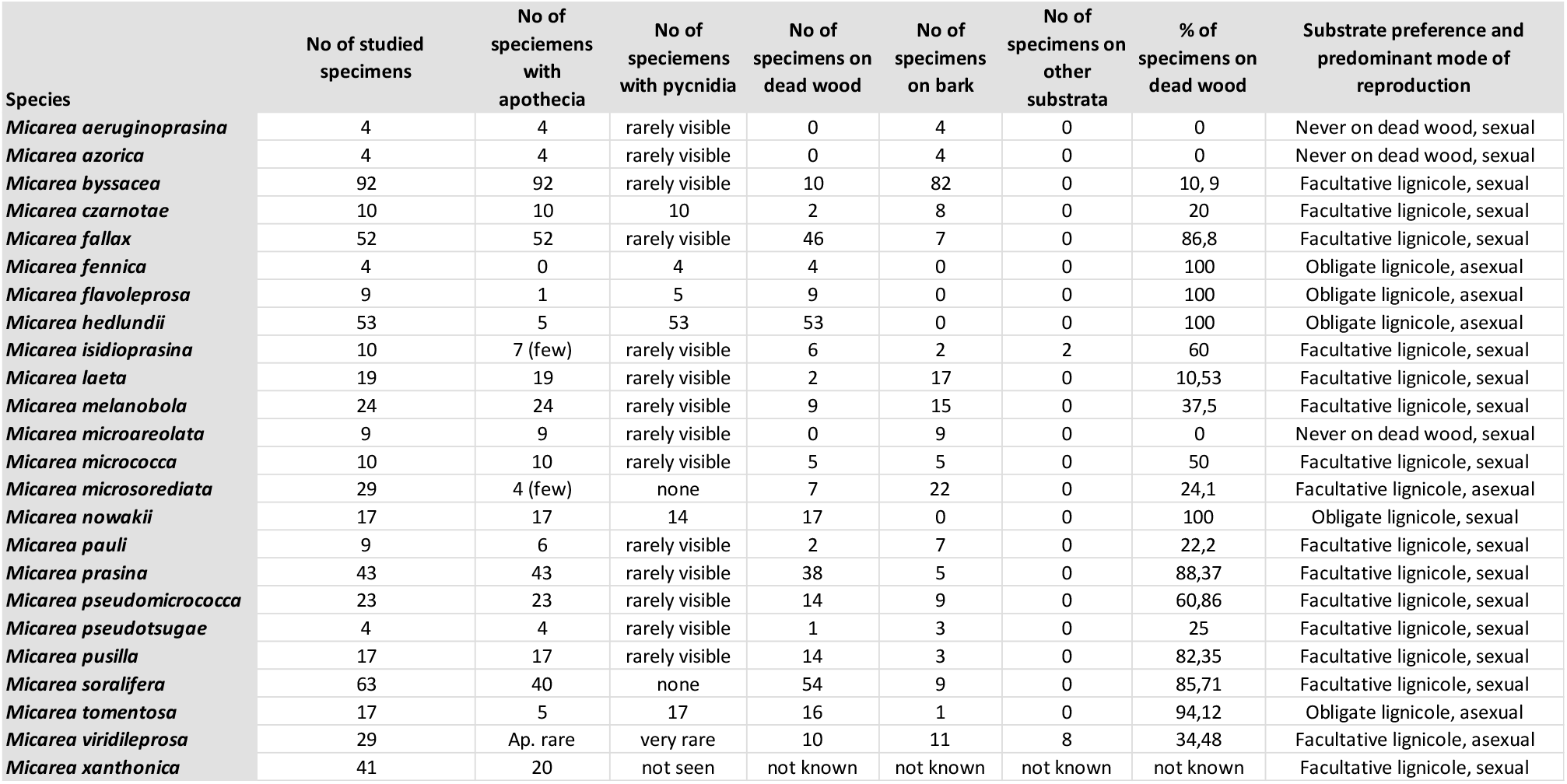
Number of studied specimens, their reproduction mode and substrate requirement. Five species are recorded based on literature: *M. herbarum* and *M. meridionalis* (van den Boom 2017), *M. subviridescens* (van den Boom 2003 & NBN Atlas online records from herbarium E), *M. viridileprosa* (van den Boom & Coppins 2001) and *M. xanthonica* (Coppins & Tønsberg 2001).

Many species in the *M. prasina* group are important colonizers of dead wood (eg. Czarnota 2007). Spribille et al. (2008) concluded that most of the obligate lichen species growing on dead wood are sexually reproducing crustose lichens. Contrary to this view, our study shows that *Micarea* species occupying bark are predominantly sexual in their reproduction mode, whereas species restricted to dead wood reproduce asexually.

Several interpretations are possible for explaining our results. Our main hypothesis involves the species’ life cycle. Species on decaying wood face a significant challenge as their substratum gradually changes and inevitably vanishes. When this happens, species need to colonize new suitable substrata. This may set a time limit, where asexual reproduction via mesoconidia, or with goniocysts acting as diaspores, is a faster and more effective strategy for successful colonization. In previous studies, asexual lichen lineages have been suggested to be more efficient and rapid at colonizing newly exposed substrates (Poelt 1963, Stofer et al. 2006, Ludwig et al. 2017). The complete decay of a log or stump can, however, take decades depending on the position of the tree (Stokland et al. 2012), and being restricted to an ephemeral substrate only leads to selection towards asexual reproduction if the generation time is long enough to effectively limit reproduction. Our results and literature show that many obligate lignicoles are restricted to certain decay stages (Coppins 1983; Czarnota 2007; Launis & Myllys 2019; Launis et al. 2019b), which shortens the time frame they have for growth and reproduction (Russell et al. 2013; Russell et al. 2014). *Micarea fennica, M. flavoleprosa, M. hedlundii*, and *M. tomentosa* mostly occur on late decay stages. *M. nowakii*, on the other hand, occupies hard wood in well-lit habitats. It is the only obligate lignicole in our data set that is predominately sexual (with additional mesoconidia nearly always present). This observation of *M. nowakii* can be explained by our hypothesis: selection would not work against sexual reproduction because the decay process is slower in the niche it favors and the time limit for growth and reproduction is not as strictly limited.

The background of the speciation process suggested above is unknown, but a possible population genetic scenario is that generalist species experience lower fitness on wood than on bark, and selection favors genotypes that are better adapted to wood. These adaptations to wood could have made the species less competitive on bark, ultimately leading to exclusive lignicoles.

In addition to our main hypothesis, the results could be explained by three alternative hypotheses, although we consider them as less likely. **Our first alternative hypothesis** is ecological: facultative lignicoles could experience lower fitness and produce less offspring on wood than on bark because of some inherent wood properties. Perhaps wood suppresses the formation of viable ascospores in

*Micarea* (even if apothecia may still be produced) and being less dependent on ascospores for reproduction would lead to selection favoring asexual reproduction. Although logical, we do not have any evidence to support this theory. **Our second alternative hypothesis** is dead wood rarity: species in the *M. prasina* group could be heterothallic, and dead wood rarity in space and time could lead to fragmented and geographically isolated populations – Zoller *et al*. (1999 b) showed that such a situation could hinder the possibility for finding a mating partner, and sexual reproduction would therefore become unnecessary and rare. Plasticity in the reproduction of heterothallic species has previously been recorded in several lichen genera (Honegger et al. 2004; Honegger & Zippler 2007; Ament-Velásquez et al. 2021). However, despite this possibly explaining the prevalence of asexual reproduction in obligate lignicoles, it only holds true if populations on wood have already lost their capacity to live on bark. In other words, the development of asexual reproduction is only possible after speciation has already occurred. Substrate specialization could still lead to asexual reproduction but not be a driver of speciation. To date, we have no evidence to support this theory, e.g. *Micarea* specimens collected in areas with high dead wood densities are not more frequently sexual. In addition, the mating systems of *Micarea* have not been examined. **Our third alternative hypothesis** is closely connected to the previous one: studies (e.g. Ament-Velásquez et al. 2021) have shown that lichens can reproduce asexually near the margins of their natural distribution while remaining sexual in central areas. This is usually because genetic diversity decreases towards margins making it difficult for heterothallic species to find mating partners. In theory, the obligate lignicoles in our data set could represent marginal populations of sexually reproducing species. So far, we have no evidence to support this theory either. We also consider it illogical that distribution would only affect obligate lignicoles, while facultative lignicoles remain sexual.

Our data includes two species that are predominately asexual but are not obligate lignicoles. *Micarea microsorediata* and *M. viridileprosa* occur on bark, wood, and on terrestrial substrates such as mosses. They rarely develop apothecia, and pycnidia have never been found from the latter. *Micarea microsorediata* mostly occurs in microhabitats where only few other lichen species co-exist, namely asexual *Lepraria* (Guzow-Krzemińska et al. 2019). *Micarea viridileprosa*, on the other hand, develops widely spreading thallus that consists of small granules called goniocysts, and it appears to be an opportunistic species that is occasionally found on ephemeral substrates and growing over mosses and other lichens (Czarnota 2007; van den Boom & Coppins 2001). Based on our results, *Micarea* species that are not obligate lignicoles are mostly sexual in their reproduction mode. However, the results on *M. microsorediata* and *M. viridileprosa* indicate that species occupying demanding low-light microhabitats or those that have opportunistic lifestyles benefit from asexual reproduction. Ecological strategies, niche requirements, and reproduction mode often correlate in other organisms (e.g. Simon et al. 2002; Silvertown 2008; Gomez-Mestre et al. 2012; Dańko et al. 2020).

## Conclusions

Our results show that asexual reproduction is an evolutionary reaction on substrate specialization. Based on our preliminary results the relationship between asexual reproduction mode and wood-inhabiting lifestyle appears to exist beyond the *M. prasina* group. For example, *M. anterior, M. misella*, and the taxa in the *M. nigella* group are mostly found on dead wood and are predominately asexual (Coppins 1983, Czarnota 2007, Myllys & Launis 2018). Future large-scale phylogenetic analyses will clarify how widespread this phenomenon is.

In general, asexual reproduction in lichens is often regarded as a deficient version of sexual reproduction, but we encourage instead to view it more as a gained ability and an advantageous strategy.

## Methods

### Taxon sampling

Altogether 516 *Micarea* specimens were studied in the herbarium collections of FR, GPR, H, UPS and few specimens were also studied in LE, O and Hb Malíček (Appendix). We included all available specimens of relevant species into our data set. The studied specimens are collected from bark and dead wood by several collectors between 1940 and 2019. Older specimens from the 1800s and early 1900s were excluded from our study because asexually reproducing lichens were rarely collected at the time. Our data set includes *Micarea* specimens from the best known and widely collected areas in the world, i.e. Fennoscandia (Finland, Norway, Sweden) and Central Europe (Belarus, Czech Republic, the Netherlands, Germany, Poland, western Russia).

Reproduction structures (apothecia/mesopycnidia) and substratum (bark/dead wood/other) were recorded for each specimen (Appendix). The predominant reproduction mode and substrate requirement (facultative/obligate lignicole) were then calculated for each species using percentages (Table 3). Species with over 94% occurrence on dead wood were regarded as obligate lignicoles, *M. tomentosa* being the only such species with less than 100% occurrence on dead wood (one specimen from Poland is collected from decaying bark).

Relevant literature on species ecology was also studied (Coppins 1983; Coppins & Tønsberg 2001; Czarnota 2007; Czarnota & Guzow-Krzemińska 2010; van den Boom et al. 2017; Guzow-Krzemińska et al. 2016 & 2019; Myllys & Launis 2018; Konoreva et al. 2019; Launis & Myllys 2019; Launis et al. 2019a, b; van den Boom et al. 2020; Kantelinen et al. 2021b; Weber et al. 2021), and our results are in line with it. Some of the specimens studied by previous authors are not included in our data set, however, because substrate and reproduction mode for the specimen is rarely reported.

### Morphology and chemistry

Each specimen in the data set was carefully studied and identified. Specimens were initially studied using dissecting microscopes. Anatomical features were then examined on hand-cut apothecial sections and squash preparations mounted in water using compound microscopes. Ascospore dimensions and other anatomical measurements were made in water and in potassium hydroxide (K). Chemical spot tests were performed under a compound microscope using sodium hypochlorite (C) and 10% (K) (Orange et al. 2010). Pigments were defined following Coppins (1983), Meyer & Printzen (2000), and Czarnota (2007). Some specimens were further studied using thin-layer chromatography (solvent C) following Culberson & Kristinsson (1970) and Orange et al. (2010). The crystalline granules of selected specimens were investigated using compound microscopes with polarization lenses.

### DNA extraction, polymerase chain reaction, and DNA sequencing

Selected specimens were sequenced and compared to sequences in GenBank to confirm species identifications. However, the majority of sequences have already been prepared for our previous studies (e.g. Launis et al. 2019b; Kantelinen et al. 2021a). Genomic DNA was extracted from 1–3 apothecia of specimens stored for a maximum of one year, using the DNeasy Blood & Tissue Kit (Qiagen, Maryland, USA) following the protocol described by Myllys et al. (2011). Polymerase chain reactions (PCRs) were prepared using PuReTaq Ready-To- Go PCR Beads (GE Healthcare, Chicago, Illinois, USA). Each 25-μl reaction volume contained 19 μl distilled water (dH2O), 1 μl of each primer (10 μM), and 4 μl extracted DNA. The primers listed below were used for PCR amplification and sequencing. For the ITS region, PCR was run under the following conditions: initial denaturation for 5 min at 95 °C followed by five cycles of 30 s at 95 °C (denaturation), 30 s at 58 °C (annealing), and 1 min at 72 °C (extension); for the remaining 40 cycles, the annealing temperature was decreased to 56 °C; the PCR program ended with a final extension for 7 min at 72 °C. The primers used were ITS1-LM (Myllys et al. 1999) and ITS4 (White et al. 1990). For the mtSSU gene, PCR was run under the following conditions: initial denaturation for 10 min at 95 °C followed by six cycles of 1 min at 95 °C (denaturation), 1 min at 62 °C (annealing), and 1 min 45 s at 72 °C (extension); for the remaining 35 cycles, the annealing temperature was decreased to 56 °C; the PCR program ended with a final extension of 10 min at 72 °C. The primers used were mrSSU1 and mrSSU3R (Zoller et al. 1999). For the *Mcm*7 gene, PCR was run under two different conditions depending on the primers selected. For the first protocol, initial denaturation for 10 min at 94 °C was followed by 38 cycles of 45 s at 94 °C (denaturation), 50 s at 55 °C (annealing), and 1 min at 72 °C (extension), with the PCR program ending with a final extension for 5 min at 72 °C. The primers used were MCM7_AL1r and MCM7_AL2f (Launis & Myllys 2019). The second protocol used an initial denaturation for 10 min at 94 °C, followed by 38 cycles of 45 s at 94 °C (denaturation), 50 s at 56 °C (annealing), and 1 min at 72 °C (extension); the PCR program ended with a final extension for 5 min at 72 °C. The primers used were x.Mcm7.f (Leavitt et al. 2011) and Mcm7.1348R (Schmitt et al. 2009). PCR products were cleaned and sequenced by Macrogen Inc. (Amsterdam, The Netherlands; www.macrogen.com).

### Phylogenetic analyses

Phylogenies comprising 29 ITS, 44 mtSSU, and 37 *Mcm*7 sequences were first aligned separately with MUSCLE v.3.8.31 (Edgar 2004) using the European Molecular Biology Laboratory, European Bioinformatics Institute’s (EMBL-EBI) freely available web server (http://www.ebi.ac.uk/Tools/msa/muscle/). The single gene trees did not show any strongly supported conflicts according to the approach of Kauff & Lutzoni (2002) (with threshold bootstrap values ≥ 75%), and the three data sets were combined into a concatenated matrix in PhyDE® (Phylogenetic Data Editor, http://www.phyde.de/index.html). Based on our previous studies (Launis et al. 2019a, b) and our preliminary phylogenetic reconstruction, *Micarea incrassata* Hedl. and *M. peliocarpa* (Anzi) Coppins & R. Sant. were used as outgroups. The hypervariable region at the end of the mtSSU and the ambiguously aligned region at the end of the ITS2 were removed from the analyses. The concatenated data set, including 44 terminals, was subjected to Bayesian inference using MrBayes (v. 3.2.7a) (Ronquist & Huelsenbeck 2003) and to maximum likelihood (ML) analysis using RAxML 8.1.15 (Stamatakis 2014). For the Bayesian analysis, substitution models were selected by having the MCMC procedure sample across models (Huelsenbeck et al. 2004). The convergence of the four parallel runs was checked after 500 000 generations using Tracer (v. 1.5)(Rambaut et al. 2018) and graphed using FigTree (v. 1.4.4). For the ML analysis, the combined data set was assigned to seven partitions: ITS1, 5.8S, ITS2, mtSSU, and each of the three codon positions of *Mcm*7. An independent GTR + G model was used for each subset, and branch lengths were assumed to be proportional across subsets. Node support was estimated with 1000 bootstrap replicates using the rapid bootstrap algorithm. The alignments are available from the Dryad Digital Repository (https://doi.org/xx).

### Ancestral state reconstruction

A binary matrix was prepared with character states given for each taxon (obligate lignicole: yes/no). Reconstructions were made with Mesquite 3.40 (Maddison & Maddison 2018) using parsimony and maximum likelihood methods.

In addition, substratum requirement and reproduction mode were studied by mapping states at the tips of the tree based on calculations in Table 3. Substratum requirement was mapped as: 1. obligate lignicole; 2. facultative lignicole; 3. neither. Predominant reproduction mode was mapped as: 1. asexual, i.e. mesopycnidia present and often numerous, apothecia rare or absent; 2. sexual, i.e. apothecia present and often abundant, mesopycnidia sometimes present (in some cases mesopycnidia may be present but invisible).

### Statistical test

A Fisher’s Exact test was performed for our data to test if the association between species’ reproduction modes and substrate preferences are significant across the studied species. The test statistic value is 0.0114, and the result is significant at p < .05 (Table 2).

## Acknowledgements

Financial support for this study was provided by the Finnish Museum of Natural History Botany Unit, Societas pro Fauna et Flora Fennica (personal fellowships for AK), Eötvös Research Grant (personal grant for PP) and the Academy of Finland (Grant 323711). We are grateful to the collectors and to herbarium staff at FR, GPR, H, O, and UPS for organizing the workspace for AK and for the loan of specimens.

## Contributions

A.K. conceived the study and designed it. Analyses were carried out by A.K. with input from C.P., P.P. and L.M. C.P facilitated access to the Germain specimens and provided important context for their use. A.K. led the interpretation of the data and writing of the manuscript, with valuable input from C.P., L.M., and P.P.

## Ethics declarations

The authors declare no competing interests.

## References

Ament-Velásquez, S.L., Tuovinen, V., Bergström, L., Spribille, T., Vanderpool, D., Nascimbene, J., Yamamoto, Y., Thor, G. & Johannesson, H. The Plot Thickens: Haploid and Triploid-Like Thalli, Hybridization, and Biased Mating Type Ratios in Letharia. Front. Fungal Biol. 2:656386. doi: 10.3389/ffunb.2021.656386 (2021).

Bowler, P.A. & Rundell, P.W. Reproductive strategies in lichens. Bot. J. Linn. Soc. 70, 325–340. (1975).

Buschbom, J. & Mueller, G.M. Testing “Species Pair” Hypotheses: Evolutionary Processes in the Lichen-Forming Species Complex Porpidia flavocoerulescens and Porpidia melinodes. Molecular Biology and Evolution 23, 574–586 (2006). https://doi.org/10.1093/molbev/msj063

Coppins, B.J. A taxonomic study of the lichen genus Micarea in Europe. Bull. Brit. Mus. (Nat. Hist.), Bot. 11, 17–214 (1983).

Coppins, B.J. & Tønsberg, T. A new xanthone-containing Micarea from Northwest Europe and the Pacific Northwest of North America. Lichenologist 33, 93–96 (2001)

Coppins, B.J. Micarea Fr. In: The Lichens of Great Britain and Ireland (eds. CW Smith et al.): 583–606. London: British Lichen Society (2009).

Culberson, C.F. & Kristinsson, H.D. A standardized method for the identification of lichen products. Journal of Chromatocraphy A 46, 85–93 (1970).

Czarnota, P. The lichen genus Micarea (Lecanorales, Ascomycota) in Poland. Polish Bot. Stud. 23, 1–190 (2007).

Czarnota, P. & Guzow-Krzemińska, B. A phylogenetic study of the Micarea prasina group shows that Micarea micrococca includes three distinct lineages. Lichenologist 42, 7–21 (2010).

Dańko, A., Schaible, R. & Dańko, M.J. Salinity Effects on Survival and Reproduction of Hydrozoan Eleutheria dichotoma. Estuaries and Coasts 43, 360–374 (2020). https://doi.org/10.1007/s12237-019-00675-2

Gaya, E., Fernández-Brime, S., Vargas, R., Lachlan, R., Gueidan, C., Ramírez-Mejía, M., Luzoni, F. The adaptive radiation of lichen-forming Teloschistaceae is associated with sunscreening pigments and bark-to-rock substrate shift. PNAS 112, 11600–11605 (2015).

Gomez-Mestre, I., Pyron, R.A. & Wiens, J.J. Phylogenetic analyses reveal unexpected patterns in the evolution of reproductive modes in frogs. Evolution 66, 3687–3700 (2012). https://Doi.Org/10.1111/J.1558-5646.2012.01715.X

Guzow-Krzemińska, B., Sérusiaux, E., van den Boom, P.P.G., Brand, A.M., Launis, A., Łubek, A. & Kukwa, M. Understanding the evolution of phenotypical characters in the Micarea prasina group (Pilocarpaceae) and descriptions of six new species within the group. MycoKeys 57, 1–30 (2019).

Huelsenbeck, J.P., Larget, B. & Alfaro, M.E. Bayesian Phylogenetic Model Selection Using Reversible Jump Markov Chain Monte Carlo. Molecular Biology and Evolution 21, 1123–1133 (2004). https://doi.org/10.1093/molbev/msh123

Kantelinen, A., Hyvärinen, M., Kirika, P. & Myllys, L. Four new Micarea species from the montane cloud forests of Taita Hills, Kenya. Lichenologist 53, 81–94 (2021a). doi:10.1017/S0024282920000511

Kantelinen, A., Westberg, M., Owe-Larsson, B. & Svensson, M. New Micarea records from Norway and Sweden and an identification key to the M. prasina group in Europe. Graphis Scripta 33, 17–28 (2021b).

Karunarathne, P., Schedler, M., Martínez, E.J., Honfi, A.I., Novichkova, A. & Hojsgaard, D. Intraspecific ecological niche divergence and reproductive shifts foster cytotype displacement and provide ecological opportunity to polyploids. Ann. Bot. 121, 1183–1196 (2018).

Kraichak, E., Divakar, P.K., Crespo, A., Leavitt, S.D., Nelsen, M.P., Lücking, R. & Lumbsch, H.T. A Tale of two Hyper-diversities: Diversification dynamics of the two largest families of lichenized fungi. Sci. Rep. DOI: 10.1038/srep100288 (2015).

Launis, A. & Myllys, L. Micarea byssacea new to North America and Micarea hedlundii new to Maine, Michigan and Quebec. Opuscula Philolichenum 13, 84–90 (2014).

Launis, A. & Myllys, L. Micarea fennica, a new lignicolous lichen species from Finland. Phytotaxa 409, 179–188 (2019).

Launis, A., Pykälä, J., van den Boom, P., Sérusiaux, E. & Myllys, L. Four new epiphytic species in the Micarea prasina group from Europe. Lichenologist 51, 7–25 (2019a).

Launis, A., Malicek, J., Svensson, M., Tsurykau, A., Sérusiaux, E. & Myllys L. Sharpening species boundaries in the Micarea prasina group, with a new circumscription of the type species M. prasina. Mycologia 111, 574–592 (2019b).

Leavitt, S.D., Lumbsch, H.T., Stenroos, S. & Clair, L.L.S. Pleistocene speciation in North American lichenized fungi and the the impact of alternative species circumscriptions and rates of molecular evolution on divergence estimates. PLoS ONE 8, e85240 (2013).

Ludwig, L.R., Summerfield, T.C., Lord, J.M. & Singh, G. Characterization of the mating-type locus (MAT) reveals a heterothallic mating system in Knightiella splachnirima. Lichenologist 49, 373–385 (2017).

Maddison, D.R. & Maddison, W.P. Mesquite: a modular system for evolutionary analysis. Version 3.40 http://mesquiteproject.org (2018).

Meyer, B. & Printzen, C. Proposal for a standardized nomenclature and characterization of insoluble lichen pigments. Lichenologist 32, 571–583 (2000).

Myllys, L., Velmala, S., Holien, H., Halonen, P., Wang, L.S. & Goward, T. Phylogeny of the genus Bryoria. The Lichenologist 43, 617–638 (2011).

Myllys, L. & Launis, A. Additions to the diversity of lichens and lichenicolous fungi living on decaying wood in Finland. Graphis Scripta 30, 78–87 (2018).

Nakov, T., Beaulieu, J. & Alverson, A. Accelerated diversification is related to life history and locomotion in a hyperdiverse lineage of microbial eukaryotes (Diatoms, Bacillariophyta). New Phytol. 219, 462–473 (2018).

Orange, A., James, P.W. & White, F.J. Microchemical methods for the identification of lichens. British Lichen Society, 101 pp (2010).

Poelt, J. Flechtenflora und Eiszeit in Europa. Phyton (Horn) 10, 206–214 (1963).

Printzen, C. & Lumbsch, H.T. Molecular evidence for the diversification of extant lichens in the Late Cretaceous and Tertiary. Molecular Phylogenetics and Evolution 17, 379–387 (2000).

Rambaut, A., Drummond, A.J., Xie, D., Baele, G. & Suchard, M.A. Posterior Summarization in Bayesian Phylogenetics Using Tracer 1.7. Syst. Biol. 67, 901–904 (2018).

Resl, P., Fernańdez-Mendoza, F., Mayrhofer, H., Spribille, T. The evolution of fungal substrate specificity in a widespread group of crustose lichens. Proc. R. Soc. B 285, 20180640 (2018). http://dx.doi.org/10.1098/rspb.2018.0640

Ronquist, F. & Huelsenbeck, J.P. MrBayes 3: Bayesian phylogenetic inference under mixed models. Bioinformatics 19, 1572–1574 (2003).

Russell, M.B., Woodall, C.W., Fraver, S. & D’Amato, A.W. Estimates of downed woody debris decay class transitions for forests across the eastern United States. Ecological Modelling 251, 22–31 (2013).

Russell, M.B., Woodall, C.W., Fraver, S., D’Amato, A.W., Domke, G.M. & Skog, K.E. Residence times and decay rates of downed woody debris biomass/carbon in eastern US Forests. Ecosystems 17, 765–777 (2014).

Schneider, K., Resl, P. & Spribille, T. Escape from the cryptic species trap: lichen evolution on both sides of a cyanobacterial acquisition event. Molecular Ecology 25, 3453–3468 (2016).

Sérusiaux, E., Brand, A.M., Motiejūnaitè, J., Orange, A. & Coppins, B.J. Lecidea doliiformis belongs to Micarea, Catillaria alba to Biatora and Biatora lignimollis occurs in Western Europe. Bryologist 113, 333–344 (2010).

Silvertown, J. The Evolutionary Maintenance of Sexual Reproduction: Evidence from the Ecological Distribution of Asexual Reproduction in Clonal Plants. International Journal of Plant Sciences 169, 157–168 (2008).

Simon, J-C., Rispe, C. & Sunnucks, P. Ecology and evolution of sex in aphids. Trends in Ecology and Evolution 17, 34–39 (2002).

Spribille, T., Thor, G., Bunnell, F.L., Goward, T. & Björk, C.R. Lichens on dead wood: species-substrate relationships in the epiphytic lichen floras of the Pacific Northwest and Fennoscandia. Ecography 31, 741–750 (2008).

Stofer, S., Bergamini, A., Aragón, G., Carvalho, P., Coppins, B.J., Davey, S., Dietrich, M.,, Farkas, E., Kärkkäinen, K., Keller, K., Lökös, L., Lommi, S., Máguas, C., Mitchell, R., Pinho, P., Rico, V.J., Truscott, A-M., Wolseley, P.A., Watt, A. & Scheidegger, C. Species richness of lichen functional groups in relation to land use intensity. Lichenologist 38, 331–353 (2006).

Stokland, J.N., Siitonen, J. & Jonsson, B.G. Biodiversity in dead wood. Cambridge University Press, Cambridge, pp. 412 (2012).

Tripp, E.A. Is asexual reproduction an evolutionary dead end in lichens? Lichenologist 48, 559–580 (2016).

Tripp, E.A. & Lendemer, J.C. Twenty-seven modes of reproduction in the obligate lichen symbiosis. Brittonia 70, 1–14 (2018).

Vamosi, J.C. & Vamosi, S.M. Factors influencing diversification in angiosperms: at the crossroads of intrinsic and extrinsic traits. Amer. J. Bot. 98, 460–471 (2011).

van den Boom, P. & Coppins, B.J. Micarea viridileprosa sp. nov., an overlooked lichen species from Western Europe. Lichenologist 33, 87–91 (2001).

Wagner, C.E., Harmon, L.J. & Seehausen, O. Ecological opportunity and sexual selection together predict adaptive radiation. Nature 487: 366–369 (2012).

Weber, L., Printzen, C., Bässler, C. & Kantelinen, A. Seven Micarea (Pilocarpaceae) species new to Germany and notes on deficiently known species in the Bavarian forest. Herzogia 34, 5–17 (2021).

White, T.J., Bruns, T., Lee, S. & Taylor, J.W. Amplification and direct sequencing of fungal ribosomal RNA genes for phylogenetics. In: Innis MA, Gelfand DH, Sninsky JJ, White TJ, eds. PCR protocols: A guide to the methods and applications. New York, NY: Academic Press. p. 315–322 (1990).

Widhelm, T.J., Bertoletti, F.R, Asztalos, M.J., Mercado-Díaz, J.A., Huang, J-P., Moncada, B., Lücking, R., Magain, N., Sérusiaux, E., Goffinet, B., Crouch, N., Mason-Gamer, R., Lumbsch, H.T. Oligocene origin and drivers for diversification in the genus Sticta (Lobariaceae, Ascomycota). Mol. Phyl. Evol. 126, 58–73 (2018).

Zoller, S., Scheidegger, C. & Sperisen, C. PCR primers for the amplification of mitochondrial small subunit ribosomal DNA of lichen-forming ascomycetes. Lichenologist 31, 511–516 (1999).

Zoller, S., Lutzoni, F. & Scheidegger, C. Genetic variation within and among populations of the threatened lichen Lobaria pulmonaria in Switzerland and implications for its conservation. Molecular Ecology 8, 2049–2059 (1999).

