## Appendix for "Lichen speciation is sparked by a substrate requirement shift and reproduction mode differentiation"

**Appendix** Studied 516 herbarium specimens in FR, GPR, H, LE, O, UPS and Hb Malíček. Reproduction structures (apothecia/mesopycnidia) and substratum (bark/dead wood/other), collector and herbarium number are marked for each specimen.

| **Species** | **Reproduction structures** | | | **Substratum** | | | **Collection country** | **Collector** | **Voucher number** | **Herbarium** |
|---|---|---|---|---|---|---|---|---|---|---|
| | Apothecia | Mesopycnidia (if visible) | Goniocysts or soralia | Wood | Bark | Other | | | | |
| *Micarea byssacea* | x | | x | | x | | Poland | Kowalewska | UGDA-L-26393 | UGDA |
| *Micarea byssacea* | x | | x | | x | | Poland | Kowalewska | UGDA-L-26354 | UGDA |
| *Micarea byssacea* | x | | x | | x | | Poland | Kukwa | 19988 | UGDA |
| *Micarea byssacea* | x | | x | | x | | Poland | Kukwa | 19210 | UGDA |
| *Micarea byssacea* | x | | x | | x | | Poland | Kukwa | 17362 | UGDA |
| *Micarea byssacea* | x | | x | | x | | Poland | Kukwa | 17031 | UGDA |
| *Micarea byssacea* | x | | x | | x | | Poland | Kowalewska | UGDA-L-23026 | UGDA |
| *Micarea byssacea* | x | | x | | x | | Poland | Kukwa | UGDA-L-22707 | UGDA |
| *Micarea byssacea* | x | | x | | x | | Poland | Kukwa | 17130 | UGDA |
| *Micarea byssacea* | x | | x | x | | | Poland | Kukwa | 15612a | UGDA |
| *Micarea byssacea* | x | | x | | x | | Poland | Kukwa | 15605 | UGDA |
| *Micarea byssacea* | x | | x | x | | | Poland | Kukwa | 15582 | UGDA |
| *Micarea byssacea* | x | | x | x | | | Poland | Kukwa | 17233 | UGDA |
| *Micarea byssacea* | x | | x | | x | | Poland | Kukwa | 14157 | UGDA |
| *Micarea byssacea* | x | | x | | x | | Poland | Kukwa | 15760 | UGDA |
| *Micarea byssacea* | x | | x | | x | | Poland | Kukwa | 15782 | UGDA |
| *Micarea byssacea* | x | | x | | x | | Poland | Kukwa | 15953 | UGDA |
| *Micarea byssacea* | x | | x | | x | | Poland | Kukwa | 12873 | UGDA |
| *Micarea byssacea* | x | | x | | x | | Poland | Kukwa | 8315 | UGDA |
| *Micarea byssacea* | x | | x | x | | | Poland | Kukwa | 12975 | UGDA |
| *Micarea byssacea* | x | | x | | x | | Poland | Kukwa | 12626 | UGDA |
| *Micarea byssacea* | x | | x | | x | | Poland | Kukwa | 12084 | UGDA |
| *Micarea byssacea* | x | | x | | x | | Poland | Kukwa | 8318 | UGDA |
| *Micarea byssacea* | x | | x | | x | | Poland | Kukwa | UGDA-L-18880 | UGDA |

| | | | | | | | | | | |
|---|---|---|---|---|---|---|---|---|---|---|
| *Micarea byssacea* | x | | x | | x | | Poland | Kukwa | 8158 | UGDA |
| *Micarea byssacea* | x | | x | | x | | Poland | Kukwa | 13519 | UGDA |
| *Micarea byssacea* | x | | x | | x | | Poland | Kukwa | UGDA-L-21303 | UGDA |
| *Micarea byssacea* | x | | x | | x | | Poland | Kukwa | 13705 | UGDA |
| *Micarea byssacea* | x | | x | | x | | Poland | Kowalewska | UGDA-L-21822 | UGDA |
| *Micarea byssacea* | x | | x | | x | | Poland | Kowalewska | UGDA-L-21386 | UGDA |
| *Micarea byssacea* | x | | x | | x | | Poland | Kwiatkowska | UGDA-L-23984 | UGDA |
| *Micarea byssacea* | x | | x | | x | | Poland | Kukwa | 17433 | UGDA |
| *Micarea byssacea* | x | | x | | x | | Poland | Kukwa | 17435 | UGDA |
| *Micarea byssacea* | x | | x | x | | | Germany | Schneider | FR-0262918 | FR |
| *Micarea byssacea* | x | x | x | | x | | Germany | Printzen & students | TS2-88-2 | FR |
| *Micarea byssacea* | x | | x | | x | | Germany | Printzen & students | TS2-88-13 | FR |
| *Micarea byssacea* | x | | x | | x | | Germany | Printzen & students | TS2-88-16-6 | FR |
| *Micarea byssacea* | x | | x | | x | | Germany | Printzen & students | TS2-99-3-1 | FR |
| *Micarea byssacea* | x | | x | x | x | | Germany | Printzen & students | TS2-88-19-16 | FR |
| *Micarea byssacea* | x | | x | x | | | Germany | Printzen & students | TS2-88-16_20 | FR |
| *Micarea byssacea* | x | | x | x | | | Germany | Printzen & students | TS2-88-16_11 | FR |
| *Micarea byssacea* | x | | x | | x | | Germany | Printzen & students | TS2-99-7_12 | FR |
| *Micarea byssacea* | x | | x | x | | | Germany | Printzen & students | TS2-99-9_15 | FR |
| *Micarea byssacea* | x | | x | | x | | Germany | Printzen & students | TS2-99-10_14 | FR |
| *Micarea byssacea* | x | | x | | x | | Germany | Printzen & students | TS2-99-10_22 | FR |
| *Micarea byssacea* | x | x | x | | x | | Sweden | Nordin, Sundin & Thor | (L-161761)365715 | UPS |
| *Micarea byssacea* | x | (x) | x | | x | | Sweden | Nordin | L-917462 | UPS |
| *Micarea byssacea* | x | | x | x | x | | Sweden | Nordin | L-891109 | UPS |
| *Micarea byssacea* | x | | x | | x | | Sweden | Westberg, Ekman, Hirschheydt | L-872309 | UPS |
| *Micarea byssacea* | x | | x | | x | | Sweden | Westberg, Ekman, Hirschheydt | L-872172 | UPS |

| | | | | | | | | | | |
|---|---|---|---|---|---|---|---|---|---|---|
| *Micarea byssacea* | x | | x | | x | | Sweden | Nordin | (L-202817)506146 | UPS |
| *Micarea byssacea* | x | | x | | x | | Sweden | Ekman, Westberg, Svensson, Hirschheydt | L-872214 | UPS |
| *Micarea byssacea* | x | | x | | x | | Sweden | Nordin | L-797414 | UPS |
| *Micarea byssacea* | x | | x | x | | | Sweden | Westberg | L-950851 | UPS |
| *Micarea byssacea* | x | | x | | x | | Finland | Pykälä | 53268 | H |
| *Micarea byssacea* | x | x | x | | x | | Finland | Pykälä | 53170 | H |
| *Micarea byssacea* | x | | x | | x | | Finland | Pykälä | 53067 | H |
| *Micarea byssacea* | x | | x | | x | | Finland | Pykälä | 53083 | H |
| *Micarea byssacea* | x | | x | | x | | Finland | Pykälä | 53074 | H |
| *Micarea byssacea* | x | | x | | x | | Finland | Pykälä | 54169 | H |
| *Micarea byssacea* | x | | x | | x | | Finland | Pykälä | 53273 | H |
| *Micarea byssacea* | x | | x | | x | | Finland | Pykälä | 53278 | H |
| *Micarea byssacea* | x | | x | | x | | Finland | Launis (Kantelinen) | 289101 | H |
| *Micarea byssacea* | x | | x | | x | | Finland | Launis (Kantelinen) | 289102 | H |
| *Micarea byssacea* | x | | x | | x | | Finland | Launis (Kantelinen) | 289104 | H |
| *Micarea byssacea* | x | | x | | x | | Finland | Launis (Kantelinen) | 208121 | H |
| *Micarea byssacea* | x | | x | | x | | Sweden | Czarnota | Czarnota s.n. | H |
| *Micarea byssacea* | x | | x | | x | | Finland | Pykälä | 47782 | H |
| *Micarea byssacea* | x | | x | | x | | Finland | Pykälä | 47789 | H |
| *Micarea byssacea* | x | | x | | x | | Finland | Pykälä | 47784 | H |
| *Micarea byssacea* | x | | x | | x | | Finland | Pykälä | 47788 | H |
| *Micarea byssacea* | x | | x | | x | | Russia | Himelbrant & Stepanchikova | H9220198 | H |
| *Micarea byssacea* | x | | x | | x | | Finland | Thor | 9445 | UPS |
| *Micarea byssacea* | x | | x | | x | | Finland | Kantelinen | 4670 | H |

| | | | | | | | | | | |
|---|---|---|---|---|---|---|---|---|---|---|
| Micarea byssacea | x | | x | | x | | Germany | Schneider | FR-0263061 | FR |
| Micarea byssacea | x | | x | | x | | Germany | Schneider | FR-0263020 | FR |
| Micarea byssacea | x | | x | | x | | Sweden | Mattsson | (L-95673)167568 | UPS |
| *Micarea byssacea* | x | | x | | x | | Germany | Schneider | FR-0263356 | FR |
| Micarea byssacea | x | | x | | x | | Germany | Schneider | FR-0263209 | FR |
| Micarea byssacea | x | | x | | x | | Sweden | Knutsson | L-720685 | UPS |
| Micarea byssacea | x | | x | | x | | Sweden | Knutsson | L-724517 | UPS |
| Micarea byssacea | x | | x | | x | | Sweden | Arup, Ekman & Knutsson | L-720671 | UPS |
| Micarea byssacea | x | | x | | x | | Sweden | Knutsson | L-724621 | UPS |
| Micarea byssacea | x | | x | | x | | Sweden | Knutsson | L-724617 | UPS |
| Micarea byssacea | x | | x | | x | | Sweden | Tibell | L-726862 | UPS |
| Micarea byssacea | x | | x | | x | | Sweden | Nordin, Sundin & Thor | (L-161767)365721 | UPS |
| Micarea byssacea | x | | x | | x | | Sweden | Nordin, Sundin & Thor | (L-161760)365714 | UPS |
| Micarea byssacea | x | | x | | x | | Sweden | Nordin, Sundin & Thor | (L-161763)365717 | UPS |
| Micarea byssacea | x | | x | | x | | Sweden | Thor | (L-174172)434195 | UPS |
| Micarea byssacea | x | | x | | x | | Sweden | Thor | L-745658 | UPS |
| Micarea byssacea | x | | x | | x | | Sweden | Svensson | (L-167556)399878 | UPS |
| *Micarea byssacea* | x | | x | | x | | Sweden | Thor | (L-174170)434193 | UPS |
| Micarea byssacea | x | | x | x | | | Germany | Schneider | FR-0263041 | FR |
| *Micarea czarnotae* | x | x | x | | x | | Finland | Launis (Kantelinen) | 109111 | H |
| *Micarea czarnotae* | x | x | x | x | | | Finland | Launis (Kantelinen) | 1010133 | H |
| *Micarea czarnotae* | x | x | x | x | | | Netherland | van den Boom | 50312 | LG |
| *Micarea czarnotae* | x | x | x | | x | | Poland | Czarnota | 3632 | GPN |
| *Micarea czarnotae* | x | x | x | | x | | Poland | Czarnota | 4179 | GPN |

| | | | | | | | | | | |
|---|---|---|---|---|---|---|---|---|---|---|
| *Micarea czarnotae* | x | x | x | | x | | Poland | Czarnota | 3179 | GPN |
| *Micarea czarnotae* | x | x | x | | x | | Poland | Czarnota | 4059 | GPN |
| *Micarea czarnotae* | x | x | x | | x | | Russia | Himelbrant & Stepanchikova | H9219944 | H |
| Micarea czarnotae | x | x | x | | x | | Sweden | Nordin, Sundin & Thor | (L-161768)365722 | UPS |
| Micarea czarnotae | x | x | x | | x | | Sweden | Nordin, Sundin & Thor | (L-161757)365711 | UPS |
| *Micarea fallax* | x | | x | x | | | Germany | Schneider | FR-0263156 | FR |
| *Micarea fallax* | x | | x | x | | | Germany | Schneider | FR-0263079 | FR |
| *Micarea fallax* | x | | x | x | | | Germany | Printzen & students | TS3-2,-15 | FR |
| *Micarea fallax* | x | | x | x | | | Germany | Printzen & students | TS3-15,-14 | FR |
| *Micarea fallax* | x | | x | x | | | Germany | Printzen & students | TS3-54,-7 | FR |
| *Micarea fallax* | x | | x | x | | | Germany | Printzen & students | TS3-17,-16 | FR |
| *Micarea fallax* | x | | x | x | | | Germany | Printzen & students | TS3-49,-9 | FR |
| *Micarea fallax* | x | | x | x | | | Germany | Printzen & students | TS3-35,-11 | FR |
| *Micarea fallax* | x | | x | x | | | Germany | Printzen & students | TS3-39,-13 | FR |
| *Micarea fallax* | x | | x | x | | | Germany | Printzen & students | TS3-41,-10 | FR |
| *Micarea fallax* | x | | x | x | | | Germany | Printzen & students | TS3-41,-11 | FR |
| *Micarea fallax* | x | | x | x | | | Germany | Printzen & students | TS3-37,-11 | FR |
| *Micarea fallax* | x | | x | x | | | Germany | Printzen & students | TS3-48,-13 | FR |
| *Micarea fallax* | x | | x | x | | | Germany | Printzen & students | TS3-48,-4 | FR |
| *Micarea fallax* | x | | x | x | | | Germany | Printzen & students | TS4-26,-3 | FR |
| *Micarea fallax* | x | | x | x | | | Germany | Printzen & students | TS4-21,-10 | FR |
| *Micarea fallax* | x | | x | x | | | Germany | Printzen & students | TS2-88-6 | FR |
| *Micarea fallax* | x | | x | x | | | Germany | Printzen & students | TS2-88-16-6 | FR |
| *Micarea fallax* | x | | x | x | | | Germany | Printzen & students | TS4-21,-11 | FR |
| *Micarea fallax* | x | | x | x | | | Germany | Printzen & students | TS4-60 | FR |
| *Micarea fallax* | x | | x | x | | | Germany | Printzen & students | TS4-40,-10 | FR |
| *Micarea fallax* | x | | x | x | | | Germany | Printzen & students | TS4-36,-6 | FR |

| | | | | | | | | | | |
|---|---|---|---|---|---|---|---|---|---|---|
| *Micarea fallax* | x | | x | x | | | Germany | Printzen & students | TS4-53,-12 | FR |
| *Micarea fallax* | x | | x | x | | | Germany | Printzen & students | TS4-5,-13 | FR |
| *Micarea fallax* | x | | x | x | | | Germany | Printzen & students | TS4-23,-6 | FR |
| *Micarea fallax* | x | | x | x | | | Germany | Printzen & students | TS4-40,-10 | FR |
| *Micarea fallax* | x | | x | x | | | Germany | Printzen & students | TS4-1,-5 | FR |
| *Micarea fallax* | x | | x | x | | | Germany | Printzen & students | TS4-1,-12 | FR |
| *Micarea fallax* | x | | x | x | | | Germany | Printzen & students | TS4-16,-11 | FR |
| *Micarea fallax* | x | | x | x | | | Germany | Printzen & students | TS4-2,-10 | FR |
| *Micarea fallax* | x | | x | x | | | Germany | Printzen & students | TS4-20,-8 | FR |
| *Micarea fallax* | x | | x | x | | | Germany | Printzen & students | TS3-57,-4 | FR |
| *Micarea fallax* | x | | x | x | | | Germany | Printzen & students | TS4-29,-11 | FR |
| *Micarea fallax* | x | | x | x | | | Germany | Printzen & students | TS4-20,-9 | FR |
| *Micarea fallax* | x | | x | x | | | Germany | Printzen & students | TS4-20,-14 | FR |
| *Micarea fallax* | x | | x | x | | | Germany | Printzen & students | TS3-28,-8 | FR |
| *Micarea fallax* | x | | x | x | | | Germany | Printzen & students | TS3-46,-15 | FR |
| *Micarea fallax* | x | | x | x | | | Germany | Printzen & students | TS4-30,-13 | FR |
| *Micarea fallax* | x | | x | | x | | Finland | Launis (Kantelinen) | 109115 | H |
| *Micarea fallax* | x | | x | | x | | Belarus | Tsurykau | 001c4 | H |
| *Micarea fallax* | x | | x | | x | | Czech | Malíček | 6127 | Hb Malíček |
| *Micarea fallax* | x | | x | x | | | Czech | Malíček | 11821 | Hb Malíček |
| *Micarea fallax* | x | | x | x | | | Czech | Malíček | 11992 | Hb Malíček |
| *Micarea fallax* | x | | x | | x | | Finland | Launis (Kantelinen) | 27122 | H |
| *Micarea fallax* | x | | x | x | | | Finland | Launis (Kantelinen) | 59132 | H |
| *Micarea fallax* | x | | x | | x | | Finland | Launis (Kantelinen) | 1710132 | H |
| *Micarea fallax* | x | | x | x | | | Finland | Launis (Kantelinen) | 1010138 | H |
| *Micarea fallax* | x | | x | x | | | Finland | Launis (Kantelinen) | 1010139 | H |

| | | | | | | | | | | |
|---|---|---|---|---|---|---|---|---|---|---|
| *Micarea fallax* | x | | x | x | | | Sweden | Svensson | 2398 | H |
| *Micarea fallax* | x | | x | x | | | Russia | Himelbrant, Konoreva & Stepanchikova | GL-19 | H |
| *Micarea fallax* | x | | x | | x | | Sweden | Ågren | 545 | UPS |
| *Micarea fallax* | x | | x | | x | | Poland | Kukwa | 13639 | UGDA |
| *Micarea fennica* | | x | x | x | | | Finland | Launis (Kantelinen) | 3220 | H |
| *Micarea fennica* | | x | x | x | | | Finland | Launis (Kantelinen) | 68 | H |
| *Micarea fennica* | | x | x | x | | | Norway | Klepsland | ? | O |
| *Micarea fennica* | | x | x | x | | | Norway | Klepsland | ? | O |
| *Micarea flavoleprosa* | | | x | x | | | Poland | Kukwa | 14048 | UGDA |
| *Micarea flavoleprosa* | x | | x | x | | | Poland | Kukwa | 14244 | UGDA |
| *Micarea flavoleprosa* | | x | x | x | | | Poland | Kukwa | 15582a | UGDA |
| *Micarea flavoleprosa* | (x) | | x | x | | | Czech | Malíček | 5098 | PRA |
| *Micarea flavoleprosa* | | | x | x | | | Austria | Berger | 32900 | Hb Berger |
| *Micarea flavoleprosa* | | | x | x | | | Austria | Berger | 33573 | Hb Berger |
| *Micarea flavoleprosa* | | | x | x | | | Czech | Malíček | 4699 | Hb Malíček |
| *Micarea flavoleprosa* | | | x | x | | | Czech | Malíček | 11823 | Hb Malíček |
| *Micarea flavoleprosa* | | | x | x | | | France | Sérusiaux | s.n. | LG |
| *Micarea flavoleprosa* | | x | x | x | | | Poland | Kukwa | 3168 | UGDA |
| *Micarea hedlundii* | | x | x | x | | | Poland | Czarnota | UGDA-L-9340 | UGDA |
| *Micarea hedlundii* | | x | x | x | | | Poland | Kukwa | 17364 | UGDA |
| *Micarea hedlundii* | | x | x | x | | | Poland | Kukwa | 17375 | UGDA |
| *Micarea hedlundii* | | x | x | x | | | Sweden | Weibull | L-739958 | UPS |
| *Micarea hedlundii* | | x | x | x | | | Sweden | Muhr | (L-08810) 23700 | UPS |
| *Micarea hedlundii* | | x | x | x | | | Sweden | Nordin | (L-64015)113111 | UPS |
| *Micarea hedlundii* | | x | x | x | | | Sweden | Muhr | L-591888 | UPS |

| | | | | | | | | | | |
|---|---|---|---|---|---|---|---|---|---|---|
| *Micarea hedlundii* | | x | x | x | | | Sweden | Hermansson | (L-173108)429990 | UPS |
| *Micarea hedlundii* | x | x | x | x | | | Sweden | Hermansson | (L-126535)243762 | UPS |
| *Micarea hedlundii* | | x | x | x | | | Sweden | Hermansson & Lundqvist | (L-134944)268147 | UPS |
| *Micarea hedlundii* | | x | x | x | | | Sweden | Hermansson | (L-102834)180173 | UPS |
| *Micarea hedlundii* | | x | x | x | | | Sweden | Hermansson | (L-88907)158452 | UPS |
| *Micarea hedlundii* | | x | x | x | | | Sweden | Hermansson | (L-93098)163181 | UPS |
| *Micarea hedlundii* | | x | x | x | | | Sweden | Hermansson | (L-167449)399397 | UPS |
| *Micarea hedlundii* | | x | x | x | | | Sweden | Hermansson | L-656364 | UPS |
| *Micarea hedlundii* | | x | x | x | | | Sweden | Hermansson | (L-173118)430000 | UPS |
| *Micarea hedlundii* | | x | x | x | | | Sweden | Hermansson | (L-173146)430028 | UPS |
| *Micarea hedlundii* | | x | x | x | | | Sweden | Hermansson | (L-107336)195363 | UPS |
| *Micarea hedlundii* | x | x | x | x | | | Sweden | Hermansson | L-564753 | UPS |
| *Micarea hedlundii* | | x | x | x | | | Sweden | Hermansson | (L-173145)430027 | UPS |
| *Micarea hedlundii* | | x | x | x | | | Sweden | Hermansson | (L-093190)163318 | UPS |
| *Micarea hedlundii* | | x | x | x | | | Sweden | Forslund & Koffman | (L-158163) 348193 | UPS |
| *Micarea hedlundii* | | x | x | x | | | Sweden | Hermansson | L-564719 | UPS |
| *Micarea hedlundii* | | x | x | x | | | Sweden | Hermansson | (L-125376)241345 | UPS |
| *Micarea hedlundii* | | x | x | x | | | Sweden | Hermansson | (L-102727)180066 | UPS |
| *Micarea hedlundii* | | x | x | x | | | Sweden | Tibell | (L-005555)15842 | UPS |
| *Micarea hedlundii* | | x | x | x | | | Sweden | Hermansson | (L-102818)180157 | UPS |
| *Micarea hedlundii* | | x | x | x | | | Sweden | Nordin | (L-64018) 113114 | UPS |

| | | | | | | | | | | |
|---|---|---|---|---|---|---|---|---|---|---|
| *Micarea hedlundii* | | x | x | x | | | Sweden | Muhr | L-602171 | UPS |
| *Micarea hedlundii* | | x | x | x | | | Sweden | Hermansson | (L-54952)91696 | UPS |
| *Micarea hedlundii* | | x | x | x | | | Finland | Pykälä | 34470 | H |
| *Micarea hedlundii* | | x | x | x | | | Finland | Launis (Kantelinen) | 67119 | H |
| *Micarea hedlundii* | | x | x | x | | | Finland | Launis (Kantelinen) | 1510131 | H |
| *Micarea hedlundii* | | x | x | x | | | Finland | Pykälä | 32808 | H |
| *Micarea hedlundii* | | x | x | x | | | Finland | Launis (Kantelinen) | 109101 | H |
| *Micarea hedlundii* | | x | x | x | | | Finland | Launis (Kantelinen) | 59132 | H |
| *Micarea hedlundii* | | x | x | x | | | Finland | Launis (Kantelinen) | 35708 | H |
| *Micarea hedlundii* | | x | x | x | | | Norway | Haugan | 11948 | O |
| *Micarea hedlundii* | | x | x | x | | | Norway | Haugan | 11949 | O |
| *Micarea hedlundii* | | x | x | x | | | Norway | Haugan | 11840 | O |
| *Micarea hedlundii* | | x | x | x | | | Finland | Pykälä | 47685 | H |
| *Micarea hedlundii* | | x | x | x | | | Finland | Pykälä | 47204 | H |
| *Micarea hedlundii* | | x | x | x | | | Finland | Pykälä | 47183 | H |
| *Micarea hedlundii* | | x | x | x | | | Finland | Pykälä | 47224 | H |
| *Micarea hedlundii* | | x | x | x | | | Finland | Pykälä | 31826 | H |
| *Micarea hedlundii* | | x | x | x | | | Finland | Pykälä | 47202 | H |
| *Micarea hedlundii* | | x | x | x | | | Russia | Himelbrant & Stepanchikova | Mets-01-10 | H |
| *Micarea hedlundii* | | x | x | x | | | Poland | Kukwa | 14341 | UGDA |
| *Micarea hedlundii* | | x | x | x | | | Poland | Kukwa | 14225 | UGDA |
| *Micarea hedlundii* | x | x | x | x | | | Poland | Kukwa | 15619 | UGDA |
| *Micarea hedlundii* | x | x | x | x | | | Poland | Kukwa | 15585 | UGDA |
| *Micarea hedlundii* | x | x | x | x | | | Poland | Kukwa | 15579 | UGDA |
| *Micarea hedlundii* | | x | x | x | | | Poland | Kukwa | 15962 | UGDA |

| | | | | | | | | | | |
|---|---|---|---|---|---|---|---|---|---|---|
| *Micarea isidioprasina* | x | | x | | x | | Poland | Kukwa | 14030 | UGDA |
| *Micarea isidioprasina* | | x | x | x | | | Poland | Kukwa | 13299 | UGDA |
| *Micarea isidioprasina* | | x | x | x | | | Poland | Kukwa | 17493 | UGDA |
| *Micarea isidioprasina* | x | | x | x | | | Poland | Kukwa | 14243 | UGDA |
| *Micarea isidioprasina* | x | | x | x | | | Poland | Kukwa | 14358 | UGDA |
| *Micarea isidioprasina* | x | | x | | x | | Poland | Kukwa | 14038 | UGDA |
| *Micarea isidioprasina* | | x | x | | | x | Poland | Kukwa | 14112 | UGDA |
| *Micarea isidioprasina* | x | | x | x | | | Poland | Kukwa | 14107 | UGDA |
| *Micarea isidioprasina* | x | | x | | | x | Poland | Kukwa | 13418 | UGDA |
| *Micarea isidioprasina* | x | | x | x | | | Poland | Kukwa | 17367a | UGDA |
| *Micarea laeta* | x | | x | | x | | Germany | Printzen & students | TS2-99-1-6 | FR |
| *Micarea laeta* | x | | x | x | | | Germany | Printzen & students | TS2-88-19-16 | FR |
| *Micarea laeta* | x | | x | | x | | Sweden | Nordin, Sundin & Thor | (L-161769)365723 | UPS |
| *Micarea laeta* | x | | x | | x | | Sweden | Nordin | (L-179116)449462 | UPS |
| *Micarea laeta* | x | | x | | x | | Finland | Launis (Kantelinen) | 59153a | H |
| *Micarea laeta* | x | | x | | x | | Finland | Launis (Kantelinen) | 1510131 | H |
| *Micarea laeta* | x | | x | | x | | Finland | Launis (Kantelinen) | 59153 | H |
| *Micarea laeta* | x | | x | | x | | Finland | Launis (Kantelinen) | 49151 | H |
| *Micarea laeta* | x | | x | | x | | Finland | Launis (Kantelinen) | 59154 | H |
| *Micarea laeta* | x | | x | | x | | Finland | Launis (Kantelinen) | 59155 | H |
| *Micarea laeta* | x | | x | | x | | Finland | Launis (Kantelinen) | 49152 | H |
| *Micarea laeta* | x | | x | | x | | Finland | Launis (Kantelinen) | 186152 | H |
| *Micarea laeta* | x | | x | | x | | Finland | Launis (Kantelinen) | 269141 | H |

| | | | | | | | | | | |
|---|---|---|---|---|---|---|---|---|---|---|
| *Micarea laeta* | x | | x | | x | | Finland | Launis (Kantelinen) | 286151 | H |
| *Micarea laeta* | x | | x | | x | | Finland | Launis (Kantelinen) | 1010133 | H |
| *Micarea laeta* | x | | x | | x | | Finland | Launis (Kantelinen) | 1010134 | H |
| *Micarea laeta* | x | | x | | x | | Finland | Launis (Kantelinen) | 1010135 | H |
| *Micarea laeta* | x | | x | | x | | Russia | Himelbrant & Stepanchikova | H9219948 | H |
| *Micarea laeta* | x | | x | x | | | Germany | Weber | FR-0267185 | FR |
| *Micarea melanobola* | x | | x | x | | | Germany | Printzen & students | TS3-13,-14 | FR |
| *Micarea melanobola* | x | | x | x | | | Germany | Printzen & students | TS3-12,-14 | FR |
| *Micarea melanobola* | x | | x | x | | | Germany | Printzen & students | TS3-13,-12 | FR |
| *Micarea melanobola* | x | | x | x | | | Germany | Printzen & students | TS3-5,-11 | FR |
| *Micarea melanobola* | x | | x | x | | | Germany | Printzen & students | TS3-3,-11 | FR |
| *Micarea melanobola* | x | | x | x | | | Germany | Printzen & students | TS3-1,-14 | FR |
| *Micarea melanobola* | x | | x | x | | | Germany | Printzen & students | TS2-88-18-9 | FR |
| *Micarea melanobola* | x | | x | | x | | Sweden | Westberg, Ekman, Hirschheydt | L-872026 | UPS |
| *Micarea melanobola* | x | | x | | x | | Sweden | Westberg, Ekman, Hirschheydt | L-872044 | UPS |
| *Micarea melanobola* | x | | x | | x | | Sweden | Westberg, Ekman, Hirschheydt | L-872146 | UPS |
| *Micarea melanobola* | x | | x | | x | | Sweden | Launis (Kantelinen) | L-949282 | UPS |
| *Micarea melanobola* | x | | x | | x | | Sweden | Hedlund | L-773781 | UPS |
| *Micarea melanobola* | x | | x | | x | | Finland | Pykälä | 54151 | H |
| *Micarea melanobola* | x | | x | x | | | Finland | Launis (Kantelinen) | 79133 | H |
| *Micarea melanobola* | x | | x | | x | | Finland | Launis (Kantelinen) | 27123 | H |

| | | | | | | | | | | |
|---|---|---|---|---|---|---|---|---|---|---|
| *Micarea melanobola* | x | | x | | x | | Finland | Launis (Kantelinen) | 11014 | H |
| *Micarea melanobola* | x | | x | | x | | Finland | Launis (Kantelinen) | 49141 | H |
| *Micarea melanobola* | x | | x | | x | | Finland | Launis (Kantelinen) | 116152 | H |
| *Micarea melanobola* | x | | x | | x | | Finland | Launis (Kantelinen) | 56151 | H |
| *Micarea melanobola* | x | | x | | x | | Finland | Launis (Kantelinen) | 39151 | H |
| *Micarea melanobola* | x | | x | | x | | Finland | Launis (Kantelinen) | 286152 | H |
| *Micarea melanobola* | x | | x | | x | | Finland | Launis (Kantelinen) | 266151 | H |
| *Micarea melanobola* | x | | x | | x | | Finland | Launis (Kantelinen) | 166151 | H |
| *Micarea melanobola* | x | | x | x | | | Poland | Kukwa | 13740 | UGDA |
| *Micarea microareolata* | x | | x | | x | | Finland | Launis (Kantelinen) | 59152 | H |
| *Micarea microareolata* | x | | x | | x | | Finland | Pykälä | 47783 | H |
| *Micarea microareolata* | x | | x | | x | | Finland | Pykälä | 47787 | H |
| *Micarea microareolata* | x | | x | | x | | Finland | Launis (Kantelinen) | 59133 | H |
| *Micarea microareolata* | x | | x | | x | | Finland | Launis (Kantelinen) | 89133 | H |
| *Micarea microareolata* | x | | x | | x | | Finland | Launis (Kantelinen) | 186151 | H |
| *Micarea microareolata* | x | | x | | x | | Finland | Pykälä | 47948 | H |
| *Micarea microareolata* | x | | x | | x | | Sweden | Launis (Kantelinen) | 148131 | H |
| *Micarea microareolata* | x | | x | | x | | Sweden | Launis (Kantelinen) | 148132 | H |
| *Micarea micrococca* | x | | x | x | | | Poland | Kukwa | 17233 | UGDA |
| *Micarea micrococca* | x | x | x | x | | | Germany | Schneider | FR-0263180 | FR |
| *Micarea micrococca* | x | | x | x | | | Germany | Printzen & students | TS2-88-4_3 | FR |

| | | | | | | | | | | |
|---|---|---|---|---|---|---|---|---|---|---|
| *Micarea micrococca* | x | | x | | x | | Finland | Launis (Kantelinen) | 299101 | H |
| *Micarea micrococca* | x | | x | | x | | Finland | Launis (Kantelinen) | 238131 | H |
| *Micarea micrococca* | x | | x | | x | | Finland | Launis (Kantelinen) | 1210131 | H |
| *Micarea micrococca* | x | | x | x | | | Finland | Launis (Kantelinen) | 1210132 | H |
| *Micarea micrococca* | x | x | x | x | | | Russia | Stepanchikova & Tagirdzhanova | KB-04-2013 | H |
| *Micarea micrococca* | x | | x | | x | | Sweden | Hermansson | 10511 | UPS |
| *Micarea micrococca* | x | | x | x | | | Poland | Kukwa | 15789 | UGDA |
| *Micarea microsorediata* | (x) | x | x | | x | | Germany | Schön | FR-0221443 | FR |
| *Micarea microsorediata* | x | | x | x | | | Poland | Kukwa | 17053 | UGDA |
| *Micarea microsorediata* | | x | x | | x | | Poland | Kukwa | 7721 | UGDA |
| *Micarea microsorediata* | | x | x | x | | | Poland | Kukwa | 13462 | UGDA |
| *Micarea microsorediata* | | x | x | | x | | Poland | Kukwa | 17001a | UGDA |
| *Micarea microsorediata* | x | | x | x | | | Poland | Kukwa | 14350 | UGDA |
| *Micarea microsorediata* | | x | x | | x | | Poland | Kukwa | 13420 | UGDA |
| *Micarea microsorediata* | (x) | | x | | x | | Poland | Kukwa | 13321 | UGDA |
| *Micarea microsorediata* | | x | x | | x | | Poland | Kukwa | 17592 | UGDA |
| *Micarea microsorediata* | x | | x | x | | | Poland | Kukwa | 17641 | UGDA |
| *Micarea microsorediata* | | x | x | x | | | Poland | Kukwa | 15789 | UGDA |
| *Micarea microsorediata* | | x | x | | x | | Poland | Kukwa | 19991 | UGDA |
| *Micarea microsorediata* | | x | x | | x | | Poland | Kukwa | 17379 | UGDA |

| | | | | | | | | | | |
|---|---|---|---|---|---|---|---|---|---|---|
| *Micarea microsorediata* | | x | x | | x | | Poland | Kukwa | 14002 | UGDA |
| *Micarea microsorediata* | | x | x | | x | | Poland | Kukwa | 14000 | UGDA |
| *Micarea microsorediata* | (x) | | x | | x | | Poland | Kukwa | 17546 | UGDA |
| *Micarea microsorediata* | | x | x | | x | | Poland | Kukwa | 17040 | UGDA |
| *Micarea microsorediata* | | x | x | x | | | Poland | Kukwa | 14012 | UGDA |
| *Micarea microsorediata* | | x | x | | x | | Poland | Kukwa | 13778 | UGDA |
| *Micarea microsorediata* | | x | x | | x | | Poland | Kukwa | 17032 | UGDA |
| *Micarea microsorediata* | | x | x | | x | | Poland | Kukwa | 14045 | UGDA |
| *Micarea microsorediata* | | x | x | x | | | Poland | Kukwa | 16994 | UGDA |
| *Micarea microsorediata* | | x | x | | x | | Poland | Kukwa | 19839 | UGDA |
| *Micarea microsorediata* | | | x | | x | | Poland | Kukwa | 19212 | UGDA |
| *Micarea microsorediata* | | x | x | | x | | Poland | Kukwa | 19849 | UGDA |
| *Micarea microsorediata* | x | | x | | x | | Poland | Kukwa | 19850 | UGDA |
| *Micarea microsorediata* | (x) | | x | | x | | Poland | Kukwa | 19219 | UGDA |
| *Micarea microsorediata* | | x | x | | x | | Poland | Kukwa | 19801 | UGDA |
| *Micarea microsorediata* | | x | x | | x | | Poland | Kukwa | 19800a | UGDA |
| *Micarea nowakii* | x | x | x | x | | | Germany | Schneider | FR-0263218 | FR |
| *Micarea nowakii* | x | | x | x | | | Germany | Printzen & students | TS3-17,-12 | FR |
| *Micarea nowakii* | x | | x | x | | | Germany | Printzen & students | TS3-25,-4 | FR |
| *Micarea nowakii* | x | | x | x | | | Germany | Printzen & students | TS4-56-11 | FR |
| *Micarea nowakii* | x | x | x | x | | | Sweden | Blomberg | (L-195403)488087 | UPS |

| | | | | | | | | | | |
|---|---|---|---|---|---|---|---|---|---|---|
| *Micarea nowakii* | x | x | x | x | | | Sweden | Johansson | L-695951 | UPS |
| *Micarea nowakii* | x | x | x | x | | | Sweden | Johansson | L-695952 | UPS |
| *Micarea nowakii* | x | x | x | x | | | Sweden | Johansson | L-695948 | UPS |
| *Micarea nowakii* | x | x | x | x | | | Sweden | Johansson | L-695950 | UPS |
| *Micarea nowakii* | x | x | x | x | | | Sweden | Johansson | L-695953 | UPS |
| *Micarea nowakii* | x | x | x | x | | | Sweden | Johansson | L-695954 | UPS |
| *Micarea nowakii* | x | x | x | x | | | Sweden | Johansson | L-695947 | UPS |
| *Micarea nowakii* | x | x | x | x | | | Sweden | Johansson | L-695956 | UPS |
| *Micarea nowakii* | x | x | x | x | | | Sweden | Johansson | L-695949 | UPS |
| *Micarea nowakii* | x | x | x | x | | | Sweden | Johansson | L-695955 | UPS |
| *Micarea nowakii* | x | x | x | x | | | Sweden | Svensson | L-532777 | UPS |
| *Micarea nowakii* | x | x | x | x | | | Finland | Launis (Kanteline n) | 684 | H |
| *Micarea pauli* | x | | x | | x | | Poland | Kukwa | 17240 | UGDA |
| *Micarea pauli* | x | | x | x | | | Poland | Kukwa | 13308 | UGDA |
| *Micarea pauli* | | x | x | | x | | Poland | Kukwa | 14101 | UGDA |
| *Micarea pauli* | x | | x | | x | x | Poland | Kukwa | 17621 | UGDA |
| *Micarea pauli* | x | | x | | x | | Poland | Kukwa | 17227 | UGDA |
| *Micarea pauli* | x | | x | | x | | Poland | Kukwa | 17544 | UGDA |
| *Micarea pauli* | | x | x | | x | | Poland | Kukwa | 13345 | UGDA |
| *Micarea pauli* | x | | x | x | | | Poland | Kukwa | 17619 | UGDA |
| *Micarea pauli* | | x | x | | x | x | Poland | Kukwa | 13194 | UGDA |
| Micarea prasina | x | | x | x | | | German y | Schneider | FR-0263054 | FR |
| *Micarea prasina* | x | | x | x | | | German y | Schneider | FR-0262859 | FR |
| *Micarea prasina* | x | | x | x | | | German y | Schneider | FR-0262908 | FR |
| *Micarea prasina* | x | | x | x | | | German y | Schneider | FR-0263057 | FR |
| *Micarea prasina* | x | | x | x | | | German y | Schneider | FR-0263189 | FR |
| *Micarea prasina* | x | | x | x | | | German y | Schneider | FR-0263200 | FR |
| *Micarea prasina* | x | | x | x | | | German y | Schneider | FR-0263122 | FR |
| *Micarea prasina* | x | | x | x | | | German y | Schneider | FR-0263077 | FR |

| | | | | | | | | | | |
|---|---|---|---|---|---|---|---|---|---|---|
| *Micarea prasina* | x | | x | x | | | Germany | Printzen & students | TS2-88-10-6 | FR |
| *Micarea prasina* | x | | x | x | | | Sweden | Knutsson | L-720672 | UPS |
| *Micarea prasina* | x | | x | x | | | Sweden | Knutsson | L-720686 | UPS |
| *Micarea prasina* | x | | x | x | | | Sweden | Tibell | L-649274 | UPS |
| *Micarea prasina* | x | | x | x | | | Sweden | Tibell | L-774437 | UPS |
| *Micarea prasina* | x | | x | x | | | Sweden | Nordin | (L-58794)99147 | UPS |
| *Micarea prasina* | x | | x | | x | | Sweden | Svensson | (L-159411)351595 | UPS |
| *Micarea prasina* | x | | x | | x | | Sweden | Hermansson | (L-107292)195319 | UPS |
| *Micarea prasina* | x | | x | | x | | Sweden | Hermansson | L-663581 | UPS |
| *Micarea prasina* | x | | x | | x | | Sweden | Hermansson | (L-128035)248851 | UPS |
| *Micarea prasina* | x | | x | x | | | Sweden | Hermansson | (L-111495)205340 | UPS |
| *Micarea prasina* | x | | x | | x | | Sweden | Hermansson | (L-125626)241595 | UPS |
| *Micarea prasina* | x | | x | x | | | Sweden | Nordin | (L-131960)262417 | UPS |
| *Micarea prasina* | x | | x | x | | | Sweden | Nordin | (L-86954)155852 | UPS |
| *Micarea prasina* | x | | x | | x | | Sweden | Thor | (L-165410)388122 | UPS |
| *Micarea prasina* | x | | x | x | | | Finland | Pykälä | 54351 | H |
| *Micarea prasina* | x | | x | x | | | Austria | Hafellner | 43132 | H |
| *Micarea prasina* | x | | x | x | | | Finland | Launis (Kantelinen) | 265101 | H |
| *Micarea prasina* | x | | x | x | | | Finland | Launis (Kantelinen) | 229101 | H |
| *Micarea prasina* | x | | x | x | | | Finland | Launis (Kantelinen) | 199105 | H |
| *Micarea prasina* | x | | x | x | | | Finland | Launis (Kantelinen) | 59131 | H |
| *Micarea prasina* | x | | x | x | | | Finland | Launis (Kantelinen) | 89131 | H |

| | | | | | | | | | | |
|---|---|---|---|---|---|---|---|---|---|---|
| *Micarea prasina* | x | | x | | x | | Finland | Launis (Kanteline n) | 89135 | H |
| *Micarea prasina* | x | | x | x | | | Russia | Kuznetsov a & Stepanchi kova | H9219946 | H |
| *Micarea prasina* | x | | x | x | | | Russia | Stepanchi kova | Tuters-11-2015 | H |
| *Micarea prasina* | x | | x | | x | | Sweden | Hermanss on | 19157 | UPS |
| *Micarea prasina* | x | | x | | x | | Sweden | Hermanss on | 16222b | UPS |
| *Micarea prasina* | x | | x | | x | | Sweden | Hermanss on | 6538 | UPS |
| *Micarea prasina* | x | | x | | x | | Sweden | Svensson | 454 | UPS |
| *Micarea prasina* | x | | x | x | | | Poland | Wilk | UGDA-L-17377 | UGDA |
| *Micarea prasina* | x | | x | x | | | Poland | Kukwa | 13733 | UGDA |
| *Micarea prasina* | x | | x | x | | | Poland | Kukwa | 13720 | UGDA |
| *Micarea prasina* | x | | x | x | | | Poland | Kukwa | 13373 | UGDA |
| *Micarea prasina* | x | | x | x | | | Poland | Kukwa | 13387 | UGDA |
| *Micarea prasina* | x | | x | x | | | Poland | Kukwa | 13408 | UGDA |
| *Micarea prasina* | x | | x | | x | | Poland | Kukwa | 13393 | UGDA |
| Micarea pseudomicro cocca | x | | x | x | | | German y | Schneider | FR-0263076 | FR |
| Micarea pseudomicro cocca | x | | x | x | | | Sweden | Forslund & Koffman | (L-158384)34 8414 | UPS |
| *Micarea pseudomicro cocca* | x | | x | x | | | German y | Printzen & students | TS3-15,-7 | FR |
| *Micarea pseudomicro cocca* | x | | x | x | | | German y | Printzen & students | TS3-15,-8 | FR |
| *Micarea pseudomicro cocca* | x | | x | x | | | German y | Printzen & students | TS3-42,-16 | FR |
| *Micarea pseudomicro cocca* | x | | x | x | | | German y | Printzen & students | TS3-44,-13 | FR |
| *Micarea pseudomicro cocca* | x | | x | x | | | German y | Printzen & students | TS3-46,-15 | FR |
| *Micarea pseudomicro cocca* | x | | x | x | | | German y | Printzen & students | TS4-20,-12 | FR |
| *Micarea pseudomicro cocca* | x | | x | x | | | German y | Printzen & students | TS4-20,-4 | FR |

| | | | | | | | | | | |
|---|---|---|---|---|---|---|---|---|---|---|
| *Micarea pseudomicrococca* | x | | x | x | | | Germany | Printzen & students | TS2-88-6 | FR |
| *Micarea pseudomicrococca* | x | | x | x | | | Germany | Printzen & students | TS2-88-14-4 | FR |
| *Micarea pseudomicrococca* | x | | x | x | | | Germany | Printzen & students | TS2-88-18-7 | FR |
| *Micarea pseudomicrococca* | x | | x | x | | | Germany | Printzen & students | TS2-88-17 | FR |
| *Micarea pseudomicrococca* | x | | x | | x | | Finland | Pykälä | 53268 | H |
| *Micarea pseudomicrococca* | x | | x | | x | | Finland | Pykälä | 53258 | H |
| *Micarea pseudomicrococca* | x | | x | | x | | Finland | Pykälä | 53278 | H |
| *Micarea pseudomicrococca* | x | | x | | x | | Finland | Launis (Kantelinen) | 59151 | H |
| *Micarea pseudomicrococca* | x | | x | x | | | Finland | Launis (Kantelinen) | 89132 | H |
| *Micarea pseudomicrococca* | x | | x | | x | | Finland | Launis (Kantelinen) | 258131 | H |
| *Micarea pseudomicrococca* | x | | x | | x | | Finland | Pykälä | 47579 | H |
| *Micarea pseudomicrococca* | x | | x | | x | | Finland | Pykälä | 47574 | H |
| *Micarea pseudomicrococca* | x | x | x | | x | | Russia | Stepanchikova & Tagirdzhanova | KB-06-2013 | H |
| *Micarea pseudomicrococca* | x | | x | x | | | Sweden | Forslund & Koffman | 790 | UPS |
| *Micarea pseudotsugae* | x | | x | | x | | Netherlands | van den Boom | holotyyppi, no number, collect date 5/5/2019 | UGDA |
| *Micarea pusilla* | x | | x | x | | | Germany | Printzen & students | TS3-34,-17 | FR |
| *Micarea pusilla* | x | | x | x | | | Germany | Printzen & students | TS3-34,-13 | FR |
| *Micarea pusilla* | x | | x | x | | | Germany | Printzen & students | TS4-20,-10 | FR |
| *Micarea pusilla* | x | | x | x | | | Finland | Launis (Kantelinen) | 101035 | H |

| | | | | | | | | | | |
|---|---|---|---|---|---|---|---|---|---|---|
| *Micarea pusilla* | x | | x | x | | | Czech | Vondrák | 14632 | PRA |
| *Micarea pusilla* | x | | x | x | | | Czech | Malíček | 9590 | Hb Malíček |
| *Micarea pusilla* | x | | x | x | | | Czech | Vondrák | 14634 | PRA |
| *Micarea pusilla* | x | | x | x | | | Czech | Vondrák | 14633 | PRA |
| *Micarea pusilla* | x | | x | x | | | Czech | Malíček | 9903 | Hb Malíček |
| *Micarea pusilla* | x | | x | x | | | Czech | Malíček | 9636 | Hb Malíček |
| *Micarea pusilla* | x | | x | x | | | Czech | Vondrák | 16643 | PRA |
| *Micarea pusilla* | x | | x | x | | | Czech | Malíček | 12017 | Hb Malíček |
| *Micarea pusilla* | x | | x | | x | | Finland | Launis (Kanteline n) | 1010136 | H |
| *Micarea pusilla* | x | | x | | x | | Finland | Launis (Kanteline n) | 1010137 | H |
| *Micarea pusilla* | x | | x | x | | | Russia | Vondrák | 14668 | PRA |
| *Micarea pusilla* | x | | x | x | | | Russia | Malíček | 10449 | Hb Malíček |
| *Micarea pusilla* | x | | x | | x | | Russia | Himelbran t & Stepanchi kova | H9220216 | H |
| *Micarea soralifera* | | x | x | x | | | German y | Schneider | FR-0262786 | FR |
| *Micarea soralifera* | x | | x | x | | | Poland | Kukwa | 13001 | UGDA |
| *Micarea soralifera* | | x | x | | x | | Poland | Kukwa | 19210b | UGDA |
| *Micarea soralifera* | | x | x | | x | | Poland | Kukwa | 15850 | UGDA |
| *Micarea soralifera* | x | | x | x | | | Poland | Kukwa | 15903 | UGDA |
| *Micarea soralifera* | | x | x | | x | | Poland | Kukwa | 17615 | UGDA |
| *Micarea soralifera* | x | x | x | x | | | Poland | Kukwa | 13000 | UGDA |
| *Micarea soralifera* | | x | x | | x | | Poland | Kukwa | 15626 | UGDA |
| *Micarea soralifera* | (x) | x | x | | x | | Poland | Kukwa | 17674 | UGDA |
| *Micarea soralifera* | x | | x | x | | | Poland | Kukwa | 17650 | UGDA |
| *Micarea soralifera* | x | x | x | x | | | Poland | Kukwa | 13469 | UGDA |
| *Micarea soralifera* | x | x | x | x | | | Poland | Kukwa | 15572 | UGDA |
| *Micarea soralifera* | (x) | x | x | x | | | Poland | Kukwa | 17202 | UGDA |
| *Micarea soralifera* | x | x | x | x | | | Poland | Kukwa | 17211 | UGDA |

| | | | | | | | | | | |
|---|---|---|---|---|---|---|---|---|---|---|
| *Micarea soralifera* | x | x | x | x | | | Poland | Kukwa | 17258 | UGDA |
| *Micarea soralifera* | x | x | x | x | | | Poland | Kukwa | 17261 | UGDA |
| *Micarea soralifera* | x | x | x | x | | | Poland | Kukwa | 12939 | UGDA |
| *Micarea soralifera* | x | x | x | x | | | Poland | Kukwa | 12663 | UGDA |
| *Micarea soralifera* | x | x | x | x | | | Poland | Kukwa | 12797 | UGDA |
| *Micarea soralifera* | x | x | x | x | | | Poland | Kukwa | 12949 | UGDA |
| *Micarea soralifera* | x | x | x | x | | | Poland | Kukwa | 12969 | UGDA |
| *Micarea soralifera* | x | x | x | x | | | Poland | Kukwa | 12999 | UGDA |
| *Micarea soralifera* | x | x | x | x | | | Poland | Kukwa | 5257 | UGDA |
| *Micarea soralifera* | | x | x | x | | | Poland | Kukwa | 1504 | UGDA |
| *Micarea soralifera* | x | x | x | x | | | Poland | Kukwa | 15624 | UGDA |
| *Micarea soralifera* | | x | x | x | | | Poland | Kukwa | 12473 | UGDA |
| *Micarea soralifera* | x | x | x | x | | | Poland | Kukwa | 12722 | UGDA |
| *Micarea soralifera* | x | x | x | x | | | Poland | Kukwa | 12863 | UGDA |
| *Micarea soralifera* | x | x | x | x | | | Poland | Kukwa | 14176 | UGDA |
| *Micarea soralifera* | x | x | x | x | | | Poland | Kukwa | 14154 | UGDA |
| *Micarea soralifera* | x | x | x | | x | | Poland | Kukwa | 15791 | UGDA |
| *Micarea soralifera* | (x) | x | x | x | | | Poland | Kukwa | 15938 | UGDA |
| *Micarea soralifera* | x | x | x | x | | | Poland | Kukwa | 15924 | UGDA |
| *Micarea soralifera* | x | x | x | x | | | Poland | Kukwa | 15900 | UGDA |
| *Micarea soralifera* | x | x | x | x | | | Poland | Kukwa | 15906 | UGDA |
| *Micarea soralifera* | x | x | x | x | | | Poland | Kukwa | 17490 | UGDA |
| *Micarea soralifera* | x | x | x | x | | | Poland | Kukwa | 17464 | UGDA |
| *Micarea soralifera* | | x | x | x | | | Poland | Kukwa | 17503 | UGDA |
| *Micarea soralifera* | x | x | x | x | | | Poland | Kukwa | 17518 | UGDA |
| *Micarea soralifera* | | x | x | x | | | Poland | Kukwa | 17491 | UGDA |
| *Micarea soralifera* | | x | x | | x | | Poland | Kukwa | 17492 | UGDA |
| *Micarea soralifera* | | x | x | x | | | Poland | Kukwa | 17488 | UGDA |
| *Micarea soralifera* | x | x | x | x | | | Poland | Kukwa | 13753 | UGDA |

| | | | | | | | | | | |
|---|---|---|---|---|---|---|---|---|---|---|
| *Micarea soralifera* | (x) | x | x | x | | | Poland | Kukwa | 13480 | UGDA |
| *Micarea soralifera* | | x | x | x | | | Poland | Kukwa | 13493 | UGDA |
| *Micarea soralifera* | x | x | x | x | | | Poland | Kukwa | 14037 | UGDA |
| *Micarea soralifera* | x | x | x | x | | | Poland | Kukwa | 13732 | UGDA |
| *Micarea soralifera* | x | x | x | x | | | Poland | Kukwa | 13774 | UGDA |
| *Micarea soralifera* | x | x | x | x | | | Poland | Kukwa | 13571 | UGDA |
| *Micarea soralifera* | x | x | x | x | | | Poland | Kukwa | 13270 | UGDA |
| *Micarea soralifera* | | x | x | x | | | Poland | Kukwa | 13221 | UGDA |
| *Micarea soralifera* | x | x | x | x | | | Poland | Kukwa | 14020 | UGDA |
| *Micarea soralifera* | | x | x | x | | | Poland | Kukwa | 13398 | UGDA |
| *Micarea soralifera* | x | x | x | x | | | Poland | Kukwa | 13959 | UGDA |
| *Micarea soralifera* | | x | x | x | | | Poland | Kukwa | 13764 | UGDA |
| *Micarea soralifera* | (x) | x | x | | x | | Poland | Kukwa | 17350 | UGDA |
| *Micarea soralifera* | | x | x | x | | | Sweden | Nordin | L-909820 | UPS |
| *Micarea soralifera* | | x | x | | x | | Sweden | Westberg, Ekman, Hirschheydt | L-872084 | UPS |
| *Micarea soralifera* | | x | x | x | | | Sweden | Westberg, Ekman, Hirschheydt | L-790652 | UPS |
| *Micarea soralifera* | x | x | x | x | | | Sweden | Nordin | L-797384 | UPS |
| *Micarea soralifera* | x | x | x | x | | | Sweden | Westberg, Ekman, Hirschheydt | L-790650 | UPS |
| *Micarea soralifera* | x | x | x | x | | | Sweden | Westberg & Johannesson | L-942546 | UPS |
| *Micarea soralifera* | | x | x | x | | | Finland | Launis (Kantelinen) | 3075 | H |
| *Micarea tomentosa* | x | x | x | x | | | Poland | Kukwa | 15963 | UGDA |
| *Micarea tomentosa* | | x | x | x | | | Germany | Schneider | FR-0263200 | FR |
| *Micarea tomentosa* | | x | x | | x | | Poland | Kukwa | 17555 | UGDA |
| *Micarea tomentosa* | | x | x | x | | | Poland | Kukwa | 5096 | UGDA |
| *Micarea tomentosa* | | x | x | x | | | Poland | Kukwa | 5122 | UGDA |

| | | | | | | | | | | |
|---|---|---|---|---|---|---|---|---|---|---|
| *Micarea tomentosa* | x | x | x | x | | | Poland | Kukwa | 13352 | UGDA |
| *Micarea tomentosa* | x | x | x | x | | | Poland | Kukwa | 14188 | UGDA |
| *Micarea tomentosa* | x | x | x | x | | | Sweden | Nordin | (L-113300)210588 | UPS |
| *Micarea tomentosa* | | x | x | x | | | Sweden | Svensson | (L-171669)411893 | UPS |
| *Micarea tomentosa* | x | x | x | x | | | Sweden | Hermansson | (L-88916)158461 | UPS |
| *Micarea tomentosa* | | x | x | x | | | Finland | Launis (Kantelinen) | 2431 | H |
| *Micarea tomentosa* | | x | x | x | | | Finland | Launis (Kantelinen) | 2435 | H |
| *Micarea tomentosa* | | x | x | x | | | Finland | Launis (Kantelinen) | 2437 | H |
| *Micarea tomentosa* | | x | x | x | | | Finland | Pykälä | 49725 | H |
| *Micarea tomentosa* | | x | x | x | | | Finland | Launis (Kantelinen) | 29151 | H |
| *Micarea tomentosa* | | x | x | x | | | Finland | Pykälä | 55572 | H |
| *Micarea tomentosa* | | x | x | x | | | Sweden | Delin | (L-101264)177765 | UPS |
| *Micarea tomentosa* | | x | x | x | | | Russia | Muchnik | L-14975 | LE |
| *Micrea viridileprosa* | | x | x | x | | | Sweden | Thor | (L-200234)497493 | UPS |
